# Optimal Design of Stochastic DNA Synthesis Protocols based on Generative Sequence Models

**DOI:** 10.1101/2021.10.28.466307

**Authors:** Eli N. Weinstein, Alan N. Amin, Will Grathwohl, Daniel Kassler, Jean Disset, Debora S. Marks

## Abstract

Generative probabilistic models of biological sequences have widespread existing and potential applications in analyzing, predicting and designing proteins, RNA and genomes. To test the predictions of such a model experimentally, the standard approach is to draw samples, and then synthesize each sample individually in the laboratory. However, often orders of magnitude more sequences can be experimentally assayed than can affordably be synthesized individually. In this article, we propose instead to use stochastic synthesis methods, such as mixed nucleotides or trimers. We describe a black-box algorithm for optimizing stochastic synthesis protocols to produce approximate samples from any target generative model. We establish theoretical bounds on the method’s performance, and validate it in simulation using held-out sequence-to-function predictors trained on real experimental data. We show that using optimized stochastic synthesis protocols in place of individual synthesis can increase the number of hits in protein engineering efforts by orders of magnitude, e.g. from zero to a thousand.

## 1 INTRODUCTION

Large-scale nucleic acid sequencing and synthesis is integral to modern biology and biomedicine, from biotechnology to epidemiology to neuroscience to agriculture to evolutionary biology and beyond. Generative probabilistic modeling offers a rigorous framework for analyzing large scale sequencing data and forming predictions of new sequences that can be synthesized in the laboratory. Generative models have been used, for instance, to infer underlying structural and functional constraints on protein evolution, to predict pathogen sequences that may emerge in the future, and to predict novel enzyme sequences with desired functional properties (Marks et al., 2011; Hopf et al., 2017; Weinstein and Marks, 2021; Russ et al., 2020). In order to assay the properties of predicted sequences and discover novel functional sequences, samples from generative models must be synthesized in the laboratory at scale. Large libraries are particularly important for protein engineering applications, where they are screened for hits with rare properties, e.g. a particular catalytic or binding activity.

Unfortunately, synthesizing large numbers of samples from generative sequence models is challenging. The standard approach, which we refer to as “Monte Carlo (MC) synthesis”, is to (1) sample from the model computationally, and then (2) synthesize each sample individually (Russ et al., 2020; Shin et al., 2021; Madani et al., 2021). In practice, however, MC synthesis is limited by cost: despite recent advances in synthesis technology, gene-length libraries typically do not exceed 10^4^ unique sequences (Kosuri and Church, 2014). Far larger libraries, on the order of 10^6^ – 10^13^, can be screened in many high-throughput assays. The set of likely sequences predicted by state-of-the-art generative models is often vastly larger still: a protein model with per-residue perplexity of 2 across sequences of length 100 predicts effectively 2^100^ ≈ 10^30^ sequences. Thus MC synthesis often will come nowhere near comprehensive exploration of a models’ predictions.

In principal, combinatorial and stochastic synthesis methods – such as error prone PCR and mixed nucleotides – offer an alternative approach capable of producing much larger numbers of unique sequences for the same cost. However, the sequences produced by these methods are random, and so it is unclear how to use stochastic synthesis to gain insight into the predictions of a given generative sequence model.

In this article, we describe an experimental design method – “variational synthesis” – that leverages stochastic DNA synthesis to overcome the limitations of MC synthesis. The basic idea is to optimize the parameters of the laboratory synthesis protocol to produce samples from a distribution close to the distribution of the target generative model. Variational synthesis is a rigorous approach to building ultra-large scale libraries based on generative sequence models, and can potentially dramatically accelerate the discovery of novel functional sequences.

## 2 METHOD

We consider an arbitrary target generative model that describes a probability distribution *p*(*x*) over sequences *x*. We are interested in assaying samples from the model experimentally. The standard method, MC synthesis, is to (1) draw samples *X*_1_,…,*X*_*N*_0__ ~ *p* i.i.d. computationally, and then (2) synthesize each sequence in the laboratory, deterministically. This approach is limited by the number of sequences *N*_0_ that can be affordably synthesized deterministically, typically on the order of 10^4^ or less for gene-length sequences.

As an alternative, we propose “variational synthesis” (Figure 1): (1) write down a probabilistic model *q_θ_* (*x*) of sequences produced by a stochastic synthesis protocol with experimental parameters *θ*, (2) minimize a divergence between *q_θ_* and *p* to find *q_θ_*\* ≈ *p*, (3) run the stochastic synthesis protocol in the laboratory, producing samples *X*_1_,…,*X*_*N*_1__ ~ *q_θ*_* i.i.d.. This approach is limited by the number of sequences *N*_1_ that can be affordably screened, where in general *N*_1_ can be orders of magnitude larger than N_0_, e.g. 10^6^ – 10^11^. The increase in samples comes at the cost of accuracy, since *q_θ*_* may not exactly match *p*.

**Figure 1:**
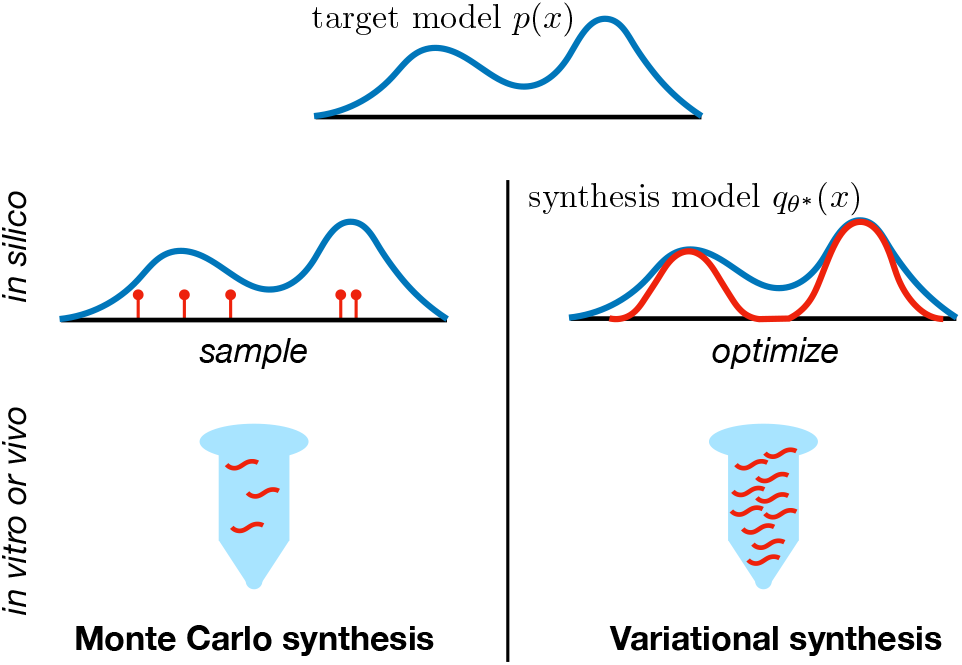
The standard synthesis approach for generative sequence models (Monte Carlo synthesis) is to sample sequences *in silico* and synthesize samples individually *in vitro.* The proposed approach (variational synthesis) is to optimize the experimental parameters of a stochastic synthesis protocol in *silico* and then run the protocol in *vitro* or in *vivo* to produce a larger number of samples.

### 2.1 Stochastic Synthesis Models

The first step of variational synthesis is to write down models *q_θ_* of stochastic synthesis protocols. We focus on five key technologies: (1) enzymatic mutagenesis, e.g. error-prone PCR or Orthorep (Wilson and Keefe, 2001; Ravikumar et al., 2018), (2) mixed nucleotide synthesis, often referred to as “degenerate codon libraries” in the context of proteins (Pazdernik and Bowersox, 2016; Mena and Daugherty, 2005), (3) mixed trimer synthesis (Kayushin et al., 1996, 2000; McMahon et al., 2018), (4) “combinatorial variant libraries” (Twist Bioscience, 2020) and (5) combinatorial assembly (Gibson et al., 2009). We focus on models of protein sequences; models of DNA or RNA are simpler.

We describe stochastic synthesis models *q_θ_* using a four-step generative process (Figure 2): (1) sample one of *M* “templates” from each of *K* “pools”, (2) join the templates together, (3) sample codons independently at each position of the combined templates and (4) translate the DNA sequence into protein. For example, consider the protocol of combinatorial assembly plus error prone PCR: we start with a library of oligos, join (assemble) a random sample of oligos into a larger sequence, and then mutagenize the sequence. Abstractly, we refer to the distribution over codons obtained by mutagenizing a particular oligo as a “template”. Techniques such as mixed nucleotides can produce alternative distributions over codons, described by different “templates”. Mathematically, let *u_kzj_*(*b*_1_, *b*_2_;, *b*_3_) denote the probability of generating codon (*b*_1_, *b*_2_, *b*_3_) at the *j*th position of template *z* in pool *k*. Let *T* be the translation matrix, defined as *T*_(*b*_1_, *b*_2_, *b*_3_)*d*_ = 1 if the codon (*b*_1_, *b*_2_, *b*_3_) codes for the amino acid *d* and *T*_(*b*_1_, *b*_2_, *b*_3_)*d*_ = 0 otherwise. (For instance, *T*_(*G, T, A)V*_ = 1 since the codon *GTA* codes for the amino acid *V*.) The complete model (Figure 2) is

**Figure 2:**
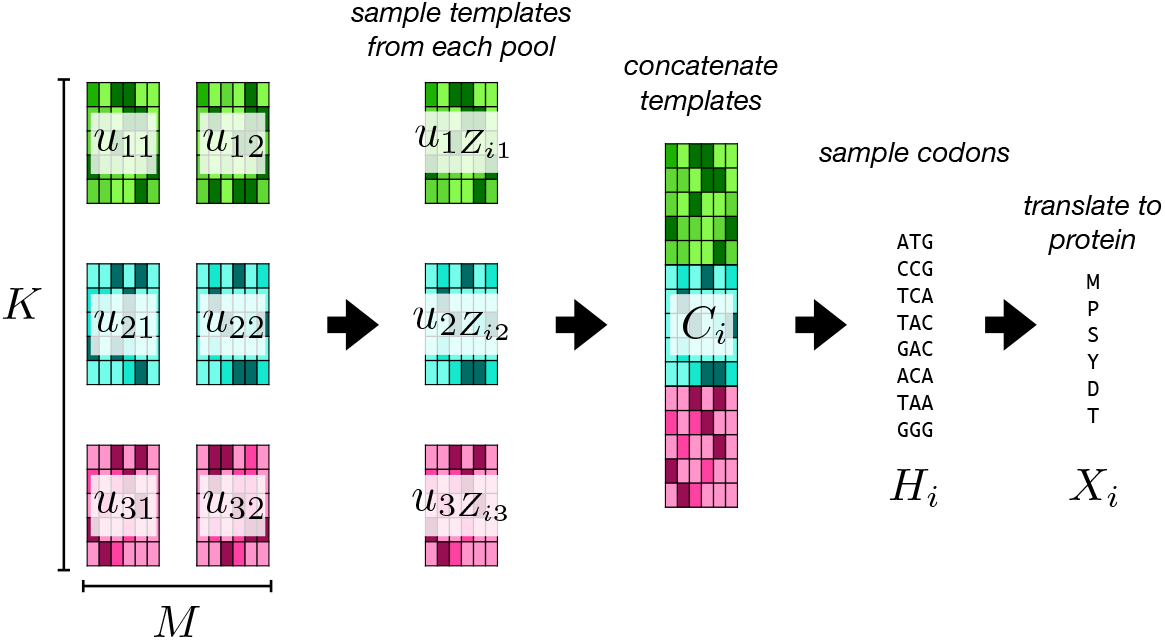
Overview of the synthesis model (Equation 1).

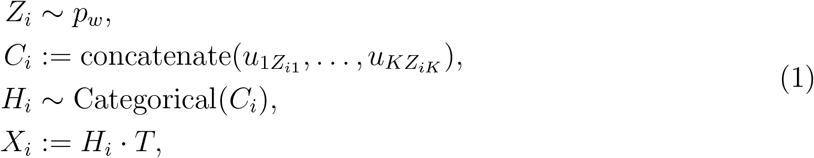

where the “concatenate” operation stacks matrices vertically, and the categorical distribution produces one-hot encoded samples based on the probabilities in each row. (Table S1 provides a notation reference.)

Different synthesis technologies impose different constraints on *p_w_*, corresponding to different assembly methods, and different constraints on *u*, corresponding to different codon diversification methods. (The biochemical basis for these different mathematical constraints is described further in Section S1.) We consider two possible constraints on *p_w_*:

1. **Fixed assembly** *Z*_*i*1_ ~ Categorical(*w*) and *Z*_*i*2_:=…:= *Z_iK_* := *Z*_*i*1_. Here we assume that there are *M* templates in each pool, and that the choice of template from the first pool dictates the choice from all the others. The experimentalist can choose the probability vector *w* ∈ Δ_*M*_, where Δ_*M*_ denotes the *M* – 1 simplex; chemically, *w* is controlled by the relative concentration of each template. In this case, the synthesis model (Equation 1) is a mixture model.
2. **Combinatorial assembly:** *Z_ik_* ~ Categorical(*w_k_*) for all *k* ∈ {1,…, *K*}. In this case each template from each pool is drawn independently. The experimentalist can choose the probability vectors *w_k_* ∈ Δ_*M*_ for each pool.

We describe constraints on the codon probabilities of each template in terms of spaces 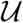 where 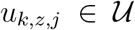. We use *v* ⊗ *v*′ to denote the outer product of two vectors *v* and *v*′. Overloading notation, for two sets of vectors *S* and *S*′, we use *S* ⊗ *S*′ to denote the set of outer products of their members, that is *S* ⊗ *S*′ := {*v* ⊗ *v*′ : *v* ∈ *S* and *v*′ ∈ *S*′}. We consider the following constraints:

1. **Arbitrary codon mixtures**: 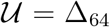. In this case, the experimentalist can choose any probability distribution over the 64 codons at each position in each template.^1^ Combinatorial variant libraries have this constraint.
2. **Finite codon mixtures**: 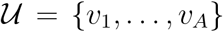 where *v_a_* ∈ Δ_64_ for all *a*. In this case, the experimentalist must first fix a library of A different codon mixtures, and then, for each position in each template, choose one of these mixtures *v_a_.* Mixed trimer synthesis protocols often have this constraint.
3. **Finite nucleotide mixtures**: 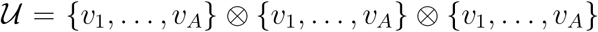 where *v_a_* ∈ Δ_4_ for all a. In this case, the experimentalist must first fix a library of A different *nucleotide* mixtures, and then, for each position in each codon in each template, choose one of these mixtures.
4. **Enzymatic mutagenesis**: 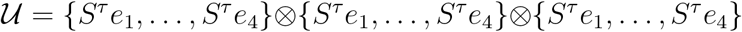 where *S* is a substitution matrix, *S^τ^* is a matrix exponential, and *e_j_* is the length 4 vector of all zeros except a one at position *j. S* has positive non-zero entries, linearly independent columns, and the sum of each column is 1. In general, the substitution matrix *S* is an intrinsic property of the chosen enzyme, while the number of rounds of mutagenesis *τ* ∈ {1,2,…} can be more easily controlled.

In summary, the parameters *θ* of the synthesis model *q_θ_* (Equation 1) that must be optimized consist of: *w* (the template probabilities), *u* (the codon probabilities), *v* (if we are using finite nucleotide or codon mixtures) and *τ* (if we are using enzymatic mutagenesis).

### 2.2 Black-Box Optimization

The second step of variational synthesis is to optimize the synthesis protocol, such that *q_θ*_* ≈ *p*. For some target/synthesis pairs – for instance, when the target is a regression model with a MuE output and fixed latent alignment (Weinstein and Marks, 2021), and the synthesis method uses fixed assembly and arbitrary codon mixtures – we can analytically and exactly match *q_θ*_* to *p* (Supplement S2.1). In most cases, however, an exact match between the target distribution and the synthesis distribution is impossible, and an analytic minimum intractable. We therefore propose an approximate optimization procedure. The primary desiderata are that it should be (1) black-box, in the sense that it can be applied to arbitrary target distributions *p* so long as *p* can be tractably sampled from, (2) scalable to large library sizes, since *q_θ_* may for instance be a mixture model with 1000 or more components and (3) able to handle large numbers of discrete parameters, since 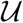 can be finite.

We propose to minimize the Kullback-Leibler (KL) divergence between the target model and the synthesis model, estimating *θ*\* := argmin_*θ*_ kl(*p*||*q_θ_*) by (1) drawing samples from the target model 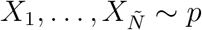 i.i.d. and (2) maximizing the log likelihood of the samples under *q_θ_* using a stochastic expectation-maximization (EM) algorithm (Cappé and Moulines, 2008). This approach only relies on samples from *p*, so can be applied whenever MC synthesis can be applied; in particular, it does not require access to likelihoods of *p*, allowing *p* to be an implicit model (e.g. a GAN). EM does not require access to derivatives of *q_θ_*(*x*) with respect to *θ*, and can easily handle categorical parameters. Finally, since the stochastic EM algorithm relies only on minibatches of data, the method is highly scalable. Sections S2.2 and S2.3 detail the algorithm and provide advice on training, including the choice of 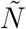.

Often the target *p* describes a distribution over variable-length protein sequences. In this case we can compute the likelihood under *q_θ_* of a protein sequence followed by a stop codon, treating the remainder of the DNA sequence as missing data (Supplement S2.4). The optimization procedure can thus be applied to *p* that produce variable-length sequences, so long as the length distribution is bounded.

## 3 RELATED WORK

Optimal design methods for stochastic synthesis have a long history, but existing techniques are in general non-probabilistic – they do not work with explicit target distributions *p* or synthesis distributions *q_θ_* – and, practically, cannot be applied to produce samples from an arbitrary generative model *p*. Methods such as LibDesign (Mena and Daugherty, 2005) and SwiftLib (Jacobs et al., 2015) optimize degenerate codon libraries to match the per-position amino acid frequencies in a multiple sequence alignment, while limiting the total size of the library. SwiftLib has for instance been used to design massive libraries of mini-protein sensors and therapeutics (Chevalier et al., 2017; Klima et al., 2021). OCoM (Parker et al., 2011) applies similar ideas to handle pairwise correlations. The recent DeCoDe method (Shimko et al., 2020) designs degenerate codon libraries to produce as many members of a set of target sequences as possible, while limiting the total size of the library; it can be interpreted probabilistically as attempting to maximize the overlap in support between a synthesis distribution *q_θ_* and a target distribution *p*, while regularizing the size of the support of *q_θ_* (Section S3.1). Meanwhile, SCHEMA and RASPP (Voigt et al., 2002; Endelman et al., 2004) are used to optimize combinatorial assembly protocols based on protein structure, and have been applied to engineer new optogenetic tools (Bedbrook et al., 2017); when the target model *p* is a Potts model that accurately reflects protein structure, variational synthesis will prefer similar solutions (Section S3.2).

Batched stochastic Bayesian optimization (Yang et al., 2019) is comparable to variational synthesis in that it is a rigorous and probabilistic approach to stochastic synthesis optimization. Unlike variational synthesis, it is focused on optimizing a reward function, rather than drawing samples from a generative sequence model. It is also not black-box, relying on the particular structure of the reward function (a Gaussian process) and focusing on just one stochastic synthesis method.

Stochastic synthesis models related to those proposed in Section 2.1 have been used in the past for inference from observational data, rather than experimental design. For instance, Tomezsko et al. (2020) use a mixture model of sequences to infer RNA structural diversity from dimethyl sulfate mutational profiling data.

Variational synthesis is inspired by variational inference (VI) (Blei et al., 2017). Both minimize a divergence between a simple approximating distribution and a target distribution (a posterior in the case of VI). Both can take advantage of the expressiveness of mixture models to achieve close matches to the target (see boosted VI by Miller et al. (2016); Guo et al. (2016); Locatello et al. (2018)). Both can be contrasted with older methods for exact sampling from a target (Markov chain Monte Carlo in the case of VI): both trade accuracy for scale, enabling large numbers of approximate samples to be drawn (computationally in the case of VI, physically in the case of variational synthesis). Both can be black-box, making sampling automatic for a large class of target distributions (Ranganath et al., 2014; Kucukelbir et al., 2017).

## 4 THEORY

### 4.1 Approximation Error

In this section, we analyze the downstream consequences of using variational synthesis in place of MC synthesis. After synthesizing (approximate) samples from *p*, the sequences will be experimentally characterized using a high-throughput assay, described by a function *f*, which provides measurements *f*(*X*_1_),…,*f*(*X_N_*) of each synthesized sequence. The assay may measure binding strength, enzymatic activity, fluorescence, etc.. *f* is assumed to be unknown before performing the experiment. We consider two distinct goals. The first goal is to estimate the average value 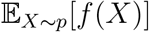. For instance, we may want to estimate the average drug resistance of future pathogen sequences predicted by *p*. Second, we may be interested in discovering a large number of sequences with a desired property, i.e. we want to maximize 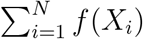 where *f*(*x*) = 1 if the sequence has the property and *f*(*x*) = 0 otherwise. E.g. if we want to engineer a new plastic-degrading protein, we want to find as many sequences as possible with high degradation rates.

#### Estimating 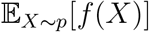

MC synthesis and variational synthesis lead to two distinct estimators for 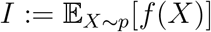, and in this section we compare their performance theoretically. In particular, the MC synthesis estimator is 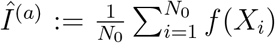 where *X*_1_,…, *X*_*N*_0__ ~ *p*, while the variational synthesis estimator is 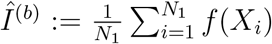 where *X*_1_,…, *X*_*N*_1__ ~ *q_θ*_*. We have no *a priori* knowledge of *f*, so to compare estimators we evaluate worst-case performance over a family of functions 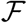. In practice, nearly all experimental assays have limited dynamic range; we therefore take 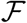 to be the set of bounded functions, 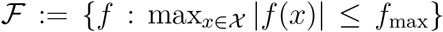, where 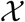 is the set of protein sequences of length less than or equal to *L.*

##### Proposition 4.1.

*The worst-case mean absolute deviation of the exact synthesis estimator satisfies*

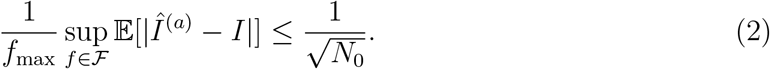

*The worst-case mean absolute deviation of the stochastic synthesis estimator satisfies*

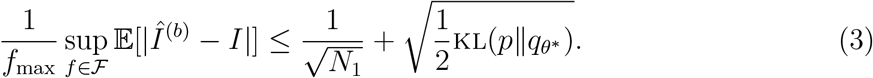

The proof, which can be found in Section S4.2, uses the integral probability metric representation of total variation along with Pinsker’s inequality. This result describes a bias-variance tradeoff: using variational synthesis in place of MC synthesis leads to less variance (since *N*_1_ > *N*_0_) but introduces bias if *q_θ*_* does not exactly match *p*. Our optimization procedure (Section 2.2) minimizes bias by minimizing kl(*p*||*q_θ_*).

If we have access to paired sequencing data, for instance if the hits of a high-throughput screen are deep-sequenced, we can remove the bias in the variational synthesis estimator via importance-weighting. We analyze this approach in Section S4.3.

#### Maximizing 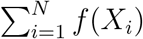

How many more hits can we expect to discover when using variational synthesis as opposed to MC synthesis? To address this question, we take *f* : *χ* → {0,1}, and compare the total number of hits when using variational synthesis, *N*_1_*Î*^(*b*)^, to the number of hits when using MC synthesis, *N*_0_*Î*^(*a*)^.

##### Corollary 4.2.

*The expected increase in hits when using variational instead of MC synthesis satisfies*

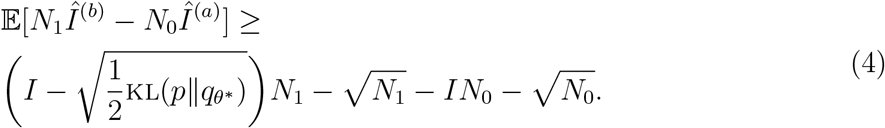

See Section S4.4 for a proof. In general *N*_1_ is much larger than *N*_0_, so the determining factor as to whether variational synthesis outperforms MC synthesis is whether *q_θ*_* is a sufficiently close approximation to *p*, i.e. whether 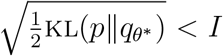. If so, the payoff from using variational synthesis can be substantial: to first order, the number of hits increases linearly with the number of sequences *N*_1_. Our optimization procedure maximizes the lower bound on the number of hits by minimizing kl(*p*||*q_θ_*).

### 4.2 Performance Limits

We have seen that the success of variational synthesis is determined by how closely *q_θ_* can match the target *p*. In this section, we analyze how closely the stochastic synthesis models described in Section 2.1 can match arbitrary target distributions *p*.

#### Limits on fixed assembly

We start by showing that synthesis protocols that use fixed assembly, and do not use enzymatic mutagenesis, can match any target distribution *p* arbitrarily well. We use *q_θ_*(*x*|*z*) as shorthand for *q_θ_* (*x*|*Z*_*i*1_ = *z*), the synthesis model distribution conditioned on the choice of template (mixture component). Let 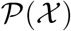 denote the set of probability distributions over *χ*. Let supp(*q_θ_*(*x*|*z*)) denote the support of the distribution *q_θ_*(*x*|*z*), i.e. the set of all *x* ∈ *χ* such that *q_θ_*(*x*|*z*) > 0.

#### Proposition 4.3.

*When using either **arbitrary codon mixtures**, **finite codon mixtures** (with A* ≥ 21), *or **finite nucleotide mixtures** (with A* ≥ *4): for any* 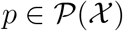 *and η* > 0 *there exists some *M* and θ such that (1)* kl(*p*||*q_θ_*) < *η and (2)* supp(*q_θ_*(*x*|*z*)) = *χ for all z* ∈ {1,…,*M*}. *When using **enzymatic mutagenesis**: there exists some* 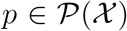 *and η* > 0 *such that for all *M* and θ, we have* kl(*p*||*q_θ_*) > η.

See Section S4.5 for a proof. The result says that as long as we are not using enzymatic mutagenesis, the target distribution *p* can be arbitrarily well approximated without resorting to individual synthesis (that is, without setting *q_θ_*(*x*|*z*) to be a delta function). Fundamentally, the problem with enzymatic mutagenesis is its discreteness: a sequence can be mutated at minimum once, so there is a minimum non-zero codon probability, given by the properties of the enzyme. This sets a limit on the “resolution” of *p* that can be matched by the synthesis procedure.^2^

#### Limits on combinatorial assembly

We next show that any synthesis protocols using combinatorial assembly cannot closely match arbitrary targets *p* even in the limit that the library size *M* goes to infinity. The result holds for any choice of 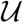.

#### Proposition 4.4.

*When using **combinatorial assembly**, so long as K* > 1, *there exists* 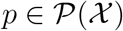 *and η* > 0 *such that for all *M* and θ, we have* kl(*p*||*q_θ_*) > *η*.

See Section S4.6 for a proof. The key problem with combinatorial assembly is that it forces templates to be independent of one another; it therefore cannot match probability distributions *p* which have correlations between regions covered by each template.

## 5 RESULTS

### 5.1 Matching Evolutionary Enzyme Models

We next evaluated the ability of variational synthesis to produce approximate samples from target protein models trained on real data. As a first target, we chose a Potts model trained on dihydrofolate reductase (DHFR) sequences from across evolution; DHFR is an enzyme crucial for nucleic acid synthesis. Potts models of protein sequences have been studied extensively, and MC synthesis from Potts models can produce functional sequences (Russ et al., 2020). We optimized each of our proposed stochastic synthesis models, setting hyperparameters based on commercially-available technologies (Section S5.2). We compared our proposed variational synthesis approach to a baseline heuristic library diversification strategy of MC synthesis plus mutagenesis: (1) draw samples from *p* and then (2) apply five rounds of mutagenesis with ePCR (Section S5.3). To evaluate how well each synthesis model matched the target distribution we estimated its per residue perplexity (Section S5.4). However, perplexity only provides a measurement of the relative quality of different synthesis procedures, rather than an absolute measurement of whether they match the data distribution. We therefore applied a Bayesian two-sample test for biological sequences – the BEAR test (Amin et al., 2021) – to determine whether *q_θ*_* in fact matches *p*, based on 100,000 samples from each (Section S5.5).

All variational synthesis methods dramatically outperform the baseline (Figure 3A), and some are capable of matching the target *p* closely, passing the two-sample test (Figure 3B). Two key determinants of the performance of the stochastic synthesis model are (1) the expressivity of the codon diversification method – that is, the size of the set of allowed 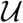 – and (2) the number of templates *M* (Section S5.2). Performance in terms of perplexity shows an improvement with increasingly large 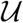 and increasing *M*. Note that due to current technology costs, when using codon mixtures, *M* must in general be small (e.g. ≤ 10) as compared to enzymatic mutagenesis or nucleotide mixtures (where *M* can be on the order of 1000). Nonetheless, using arbitrary codon mixtures with *M* = 1 templates outperforms the alternative technologies with *M* = 1000 templates.

**Figure 3:**
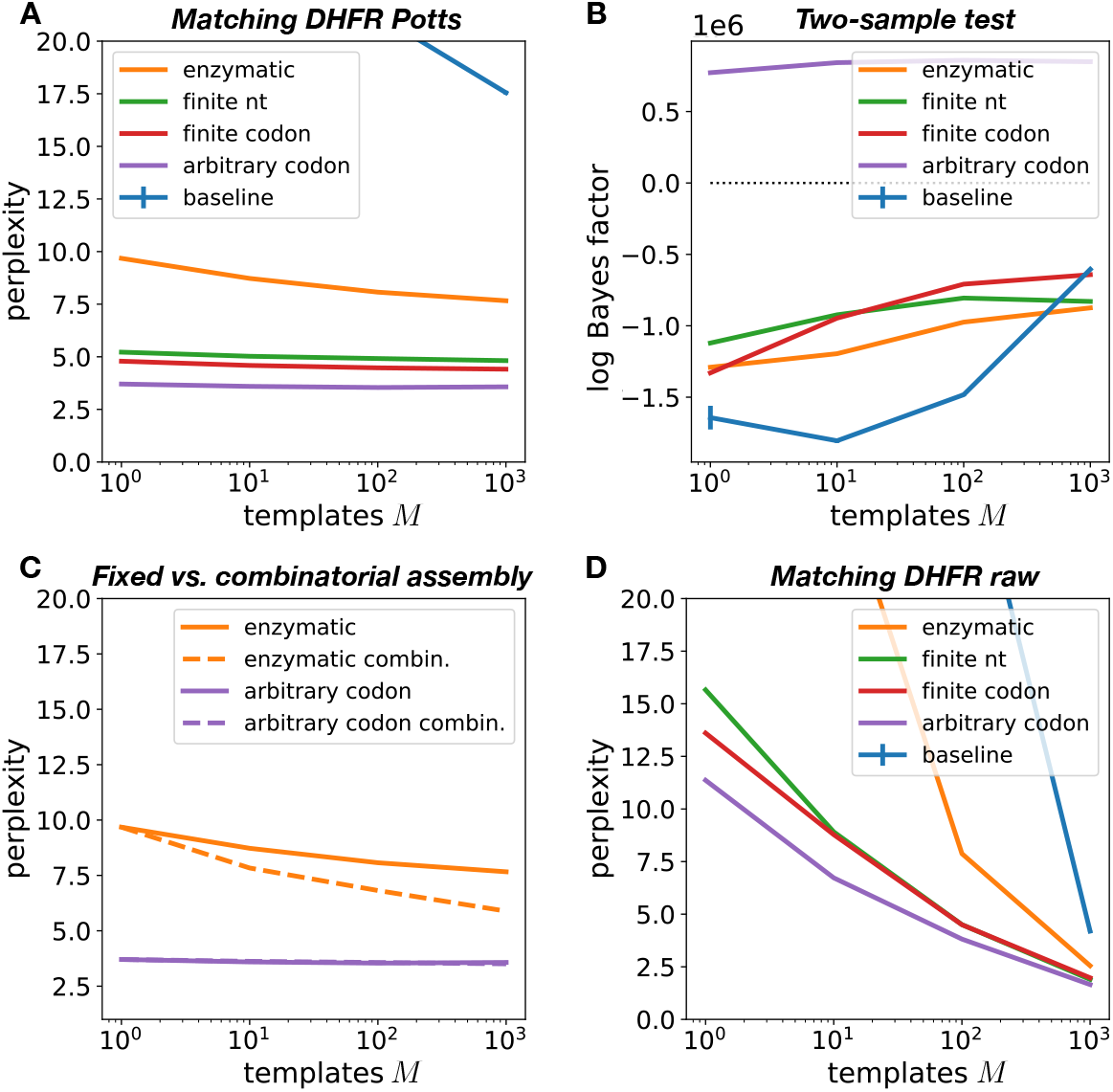
Perplexity (A) and two-sample test Bayes factor (B) of different codon diversification methods, with fixed assembly, applied to a target Potts DHFR model. Positive Bayes factors support the hypothesis that the synthesis and target distributions match. (C) Perplexity of combinatorial versus fixed assembly, applied to Potts DHFR model. (D) Perplexity of synthesis models with fixed assembly applied to unaligned DHFR sequences. Error estimates for each plot are described in detail in Section S5.7.

The advantages of combinatorial assembly over fixed assembly vary depending on the codon diversification technology. Combinatorial assembly improves perplexity when using enzymatic mutagenesis, but has little effect when using arbitrary codon mixtures (Figure 3C and Figure S3), while introducing error in the covariance matrix of *q_θ*_* (Figure S4).

We next explored the application of variational synthesis to target distributions over variable-length sequences (the DHFR Potts model was trained on aligned sequences and generates fixed-length sequences). We optimized synthesis models directly on the same evolutionary data used to train the DHFR Potts model (with gaps removed); the target here is the true evolutionary data-generating process, and unknown (Section S5.1.1). Enzymatic mutagenesis with large *M* outperforms arbitrary codon mixtures with small *M* in this case (Figures 3D and S5). The best synthesis technology can thus depend on the target.

**Figure 4:**
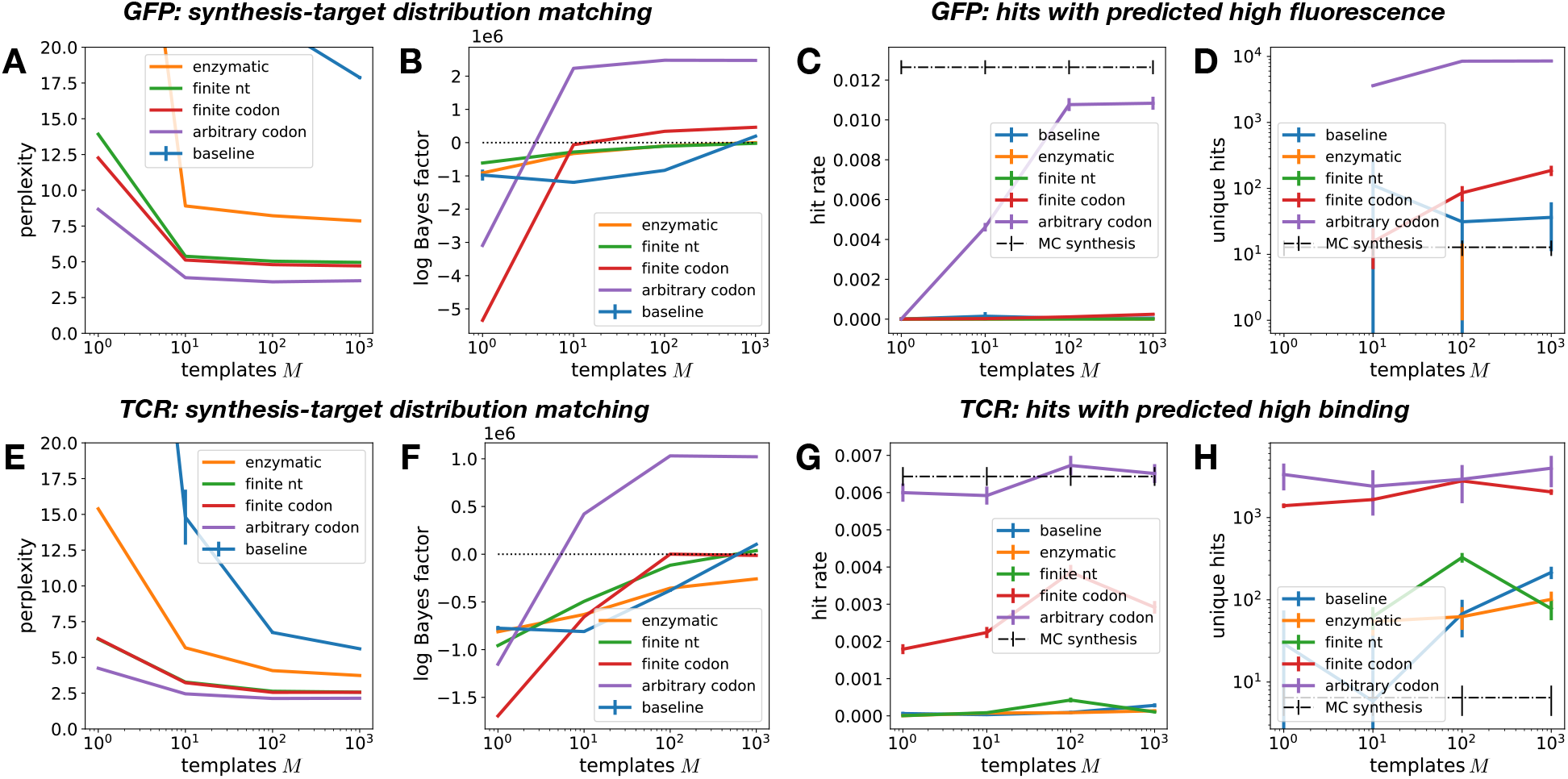
Perplexity (A) and two-sample test Bayes factor (B) for different synthesis methods applied to a target GFP model. (C) Hit rate for discovering functional sequences. (D) Expected number of unique hits in a *N*_1_ = 10^6^ library for variational synthesis, as compared to MC synthesis with a *N*_0_ = 10^3^ library (Section S5.6.4). (E-H) Same as (A-D) for a target TCR model. Error estimates for each plot are described in detail in Section S5.7.

### 5.2 Synthesizing Fluorescent Proteins

Next we sought to determine if variational synthesis can increase the number of discoveries in downstream assays, as compared to MC synthesis. To simulate the results of realistic experimental assays, we used sequence-to-function predictors trained on large-scale experimental studies. We started with green fluorescent protein (GFP), predicting fluorescence using a transformer-based semi-supervised method trained on a GFP deep mutational scan dataset and evolutionary protein data (Sarkisyan et al., 2016; Rao et al., 2019). We classified as hits sequences with predicted fluorescence above the functionality threshold specified by Sarkisyan et al. (2016) (Section S5.6.1). To construct a target *p*, we trained an unsupervised sequence model – an ICA model with MuE output, proposed in Weinstein and Marks (2021) – on evolutionarily related GFP sequences, and then fixed the latent alignment variable of the MuE to generate sequences (Section S5.1.2). Using a fixed latent alignment ensures that the fluorescence predictor, which was only trained on fixed-length sequences, can be confidently applied. Note that the fluorescence predictor was not used to construct *p* itself, so we can fairly evaluate variational synthesis in the setting where the experimental results are not known ahead of time. In general, the fluorescence predictions are quite sensitive to the input sequence – a single amino acid change can abolish fluorescence – so generating new fluorescent sequences is nontrivial (Figure S6). Only 1.3% of sequences sampled from *p* are hits, with fluorescence above the threshold specified by Sarkisyan et al. (2016).

Stochastic synthesis models with arbitrary codon mixtures and fixed assembly have low perplexities, and can pass the two-sample test with large Bayes factors at *M* ≥ 10; other methods struggle, including the baseline method (Figure 4AB). Samples from arbitrary codon models at *M* = 10 show average fluorescence similar to *p* (Figure S8), and the fraction of samples that are hits is only about half that of MC synthesis, 0.5% (Figure 4C). Meanwhile, alternative stochastic synthesis methods show hit rates below 0.05%.

Variational synthesis leads to a decrease in hit rate relative to MC synthesis, but this can be more than compensated for by the increase in the number of synthesized samples. If, for instance, *N*_1_ = 10^6^ sequences generated via variational synthesis are assayed, as opposed to *N*_0_ = 10^3^ sequences generated via MC synthesis, an estimated 3600 unique functional sequences will be discovered using variational synthesis as opposed to 10 for MC synthesis (Figure 4D; Section S5.6.4). Variational synthesis can thus provide orders-of-magnitude increases in the number of hits in protein engineering applications, with the number of hits increasing with larger values of *N*_1_ and/or *M*.

### 5.3 Synthesizing Antigen-Binding Proteins

Next we sought to evaluate the advantages of variational synthesis over MC synthesis in an application area important for human health. Understanding T cell receptor (TCR) sequences and their binding properties is crucial for understanding the immune response to infection or cancer, and engineering new TCRs with desired binding properties is crucial for immunotherapies (June et al., 2018). We trained a model of TCR sequences from a healthy donor – an ICA model with MuE output – and fixed the latent alignment variable in the MuE to define *p* (Section S5.1.3). As a held-out sequence-to-function predictor, we used Tcellmatch (Fischer et al., 2020) to predict binding to an influenza epitope (Section S5.6.2). The predictor is highly sensitive to the input sequence – a single amino acid change can abolish binding – making this a challenging problem for variational synthesis (Figure S10). Only 0.6% of samples from the target *p* are hits.

Synthesis models with arbitrary codon mixtures and fixed assembly achieve low perplexities and can pass the two-sample test with large Bayes factors (Figure 4EF). Variational synthesis with this model achieves hit rates similar to MC synthesis (Figure 4G). MC synthesis with *N*_0_ = 10^3^ generates just 6 hits on average across independent libraries; given stochasticity, it is not unlikely to see no hits at all in a given library. Variational synthesis with *N*_1_ = 10^6^ and *M* = 10 generates an expected 2400 unique hits (Figure 4H). These results suggest that variational synthesis could dramatically accelerate the discovery of new TCRs that bind specific antigens, relying only on unsupervised sequence models and not large-scale supervised sequence-to-function training data.

Close matches between *q_θ*_* and *p* turn out to be unnecessary for reaching high hit rates in this example. When using arbitrary codon mixtures or finite codon mixtures with *M* = 1, or even using finite nucleotide mixtures with *M* = 100, the two-sample test detects significant differences between *q_θ*_* and *p* (Figure 4F), but nonetheless variational synthesis achieves substantially more hits than MC synthesis (Figure 4H).

## 6 DISCUSSION

Variational synthesis trades accuracy for scale, producing large numbers of approximate samples from a target model rather than small numbers of exact samples, as in MC synthesis. When accuracy is high enough – when *q_θ*_* is sufficiently close to *p* – the payoff can be enormous, as the number of hits increases linearly with the number of assayed sequences *N*_1_. Given that many high-throughput screens can reach 10^4^ sequences or more, while individual gene synthesis rarely goes beyond *N*_0_ = 10^4^, using variational synthesis may make the difference between zero hits and a million.

The key limitations of our variational synthesis methods – and opportunities for future work – stem from the challenges of matching synthesis and target distributions. First, our synthesis models (Section 2.1) are idealizations based on manufacturers’ descriptions of the distribution of sequences their methods produce, but do not take into account possible errors, biases or limitations in the real procedure (Section S1). Developing more accurate *q_θ_* models, based on e.g. deep sequencing data, may be an important area for future work. Second, our methods for judging whether *q_θ*_* is sufficiently close to *p* are limited. Empirically, while the BEAR two-sample test appears to be excellent at distinguishing among good and bad fixed assembly models in the examples we studied, it struggles to detect the errors caused by combinatorial assembly, even when they are large enough to abolish function (Figure S9). Theoretically, tighter bounds than that in Proposition 4.1 can be proved with total variation or Wasserstein distance in place of KL, but optimizing these alternative divergences directly is a challenge (Section S4.2). For sequence-to-function predictors to be more reliable in evaluating variational synthesis methods, they must be robust to covariate shift, since switching from *p* to *q_θ*_* is, precisely, a covariate shift. Third, while our black-box optimization method allows for arbitrary target distributions *p*, it may be more effective in many cases to work with *p* for which an exactly matching *q_θ*_* can be found analytically (Section S2.1). Recent progress on mixture models as a competitor to deep generative neural network models make this approach especially promising (Richardson and Weiss, 2018).

Variational synthesis changes the calculus of what makes a successful generative sequence model and what makes a successful synthesis technology. If just 1% of the sequences sampled from an initial model A were functional, and 50% of sequences sampled from a proposed model B were functional, model B would be considered a major advance; however, if we could accurately match a stochastic synthesis protocol to model A and not to model B, then model A could easily lead to orders of magnitude more hits in practice. Meanwhile, the traditional goal of the DNA synthesis community has been large-scale individual synthesis. From a probabilistic perspective, however, it hardly makes sense to focus exclusively on methods to sample from mixtures of point masses. The recent development of methods to synthesize samples from mixture models with no constraints on codon probabilities (e.g. combinatorial variant libraries) represents a major advance outside the traditional paradigm.

Variational synthesis bridges the gap between generative sequence models and stochastic synthesis technologies, providing a rigorous approach to experimental design. We are optimistic that it will help translate powerful new generative sequence models into laboratory discoveries.

## Acknowledgments

We wish to thank Chris Sander and members of the Marks lab for valuable discussions.

## Supplementary Materials

**Table S1:**
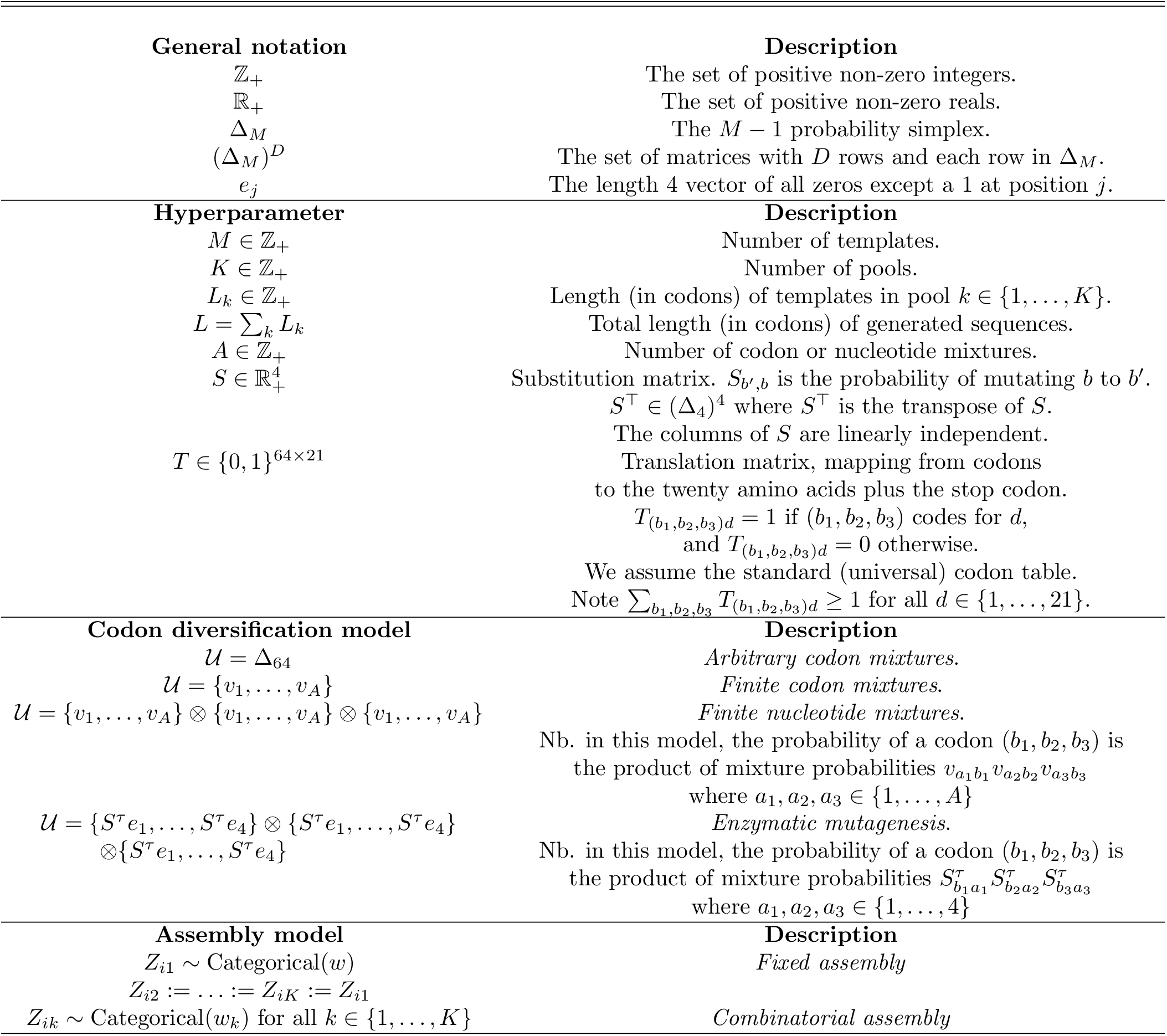
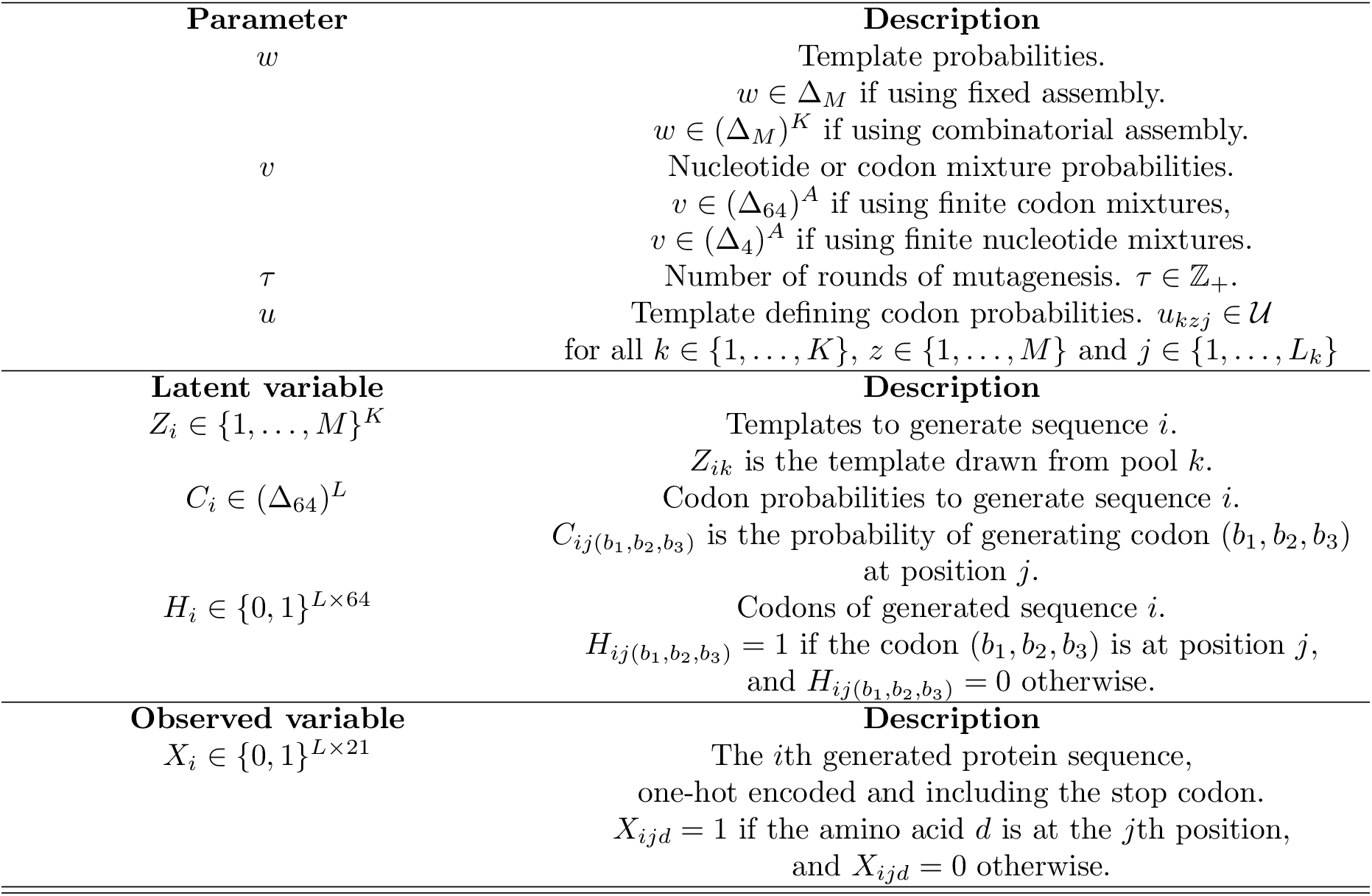
Synthesis model notation.

### S1 MODEL DETAILS AND LIMITATIONS

In this section we explain further the synthesis models proposed in Section 2.1, as well some of the limitations of our mathematical idealization.

Physically, for the finite codon or nucleotide mixture models, codon diversification happens during chemical synthesis of oligos (DNA segments). DNA in each well (or isolated reaction volume) is synthesized position by position, with mixtures of nucleotides or codons (trinucleotides) added in defined ratios one at a time, such that a large number of different molecules is eventually constructed. Twist Bioscience’s combinatorial variant libraries, which can achieve arbitrary codon mixtures, rely on proprietary technology; however, it produces analogous results (Twist Bioscience, 2020). For all of these technologies, what we refer to as a “template” corresponds physically to a very large number of molecules in an individual well, with independent nucleotide or codon probabilities at each site. We assume that the number of molecules is effectively infinite in comparison to *N*_1_, such that we do not need to account for sampling noise at this stage. We ignore the possibility of skipped positions, where nucleotides or codons randomly fail to add to the growing oligos, a type of error that is sometimes of particular concern for trimer-based synthesis. We enforce the constraint that the number of mixtures *A* is finite and small, since to the best of our knowledge commercially available technologies have this requirement, but it is not necessarily a fundamental technological constraint (Pazdernik and Bowersox, 2016).

Physically, for the enzymatic mutagenesis model, template oligos are synthesized deterministically, such that there is a large population of identical molecules in each well. Codon diversification occurs only after assembly (i.e. after oligos from different wells are combined) and may take place either *in vitro* or *in vivo.* We assume that there is an error correction mechanism after each round of mutagenesis, such that each strand of each DNA molecule has effectively gone through the same number of rounds of mutagenesis; in some ePCR protocols error correction is not used, and so alternative models may be more appropriate (Moore and Maranas, 2000; Pritchard et al., 2005). We also assume that the mutation probability depends only on individual nucleotides, and not their sequence context, although empirically dependencies on sequence context (especially the adjacent two nucleotides) can be found (Alexandrov and Stratton, 2014). Finally, we require that each template undergoes the same number of rounds of mutagenesis *τ*, with the same enzyme and thus the same *S*. For small *M*, it can be experimentally tractable in many cases to use different *τ*, and even different *S*, for each template, in which case the model should be adjusted to make *τ* and *S* depend on the template.

Physically, assembly requires joining oligos together using e.g. Gibson assembly (Gibson et al., 2009). For the fixed assembly model, the oligos corresponding to the kth template in each pool must be joined in an isolated reaction, for all *k* ∈ {1,…, *K*}; in combinatorial assembly, the sets of oligos corresponding to each template in each pool are first mixed, and then oligos from these combined pools are joined. Assembly requires short overhangs, sequences that closely match one another, at the ends of each oligo that are to be joined. Our synthesis model ignores any restrictions that come from overhangs needing to match, as well as variation in assembly probability that depend on overhang mismatch. Our model also assumes full control over the relative concentration of templates, *w.* While this is tractable for low *M*, it may be more challenging for large *M*, particularly if technologies like Dropsynth are used for fixed assembly (Plesa et al., 2018).

### S2 OPTIMIZATION DETAILS

#### S2.1 Exact Solutions

As an example of a target sequence model which we can exactly match, consider a Regress-MuE (Weinstein and Marks, 2021), which has been used for forecasting the evolution of influenza. Let **B** be a covariate vector (e.g. a future time), let **Θ** be the regression coefficients, and let **W** be the latent alignment. The predictive distribution *p*(*x*|**B**, **W**, **Θ**) can be written as Categorical(**U**), where **U** is a matrix of independent amino acid probabilities over *L* positions. We can exactly match this distribution with a synthesis model using *M* =1 templates, fixed assembly and arbitrary codon mixtures.

We can also approximate the posterior predictive distribution. Let 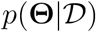 be the posterior distribution over regression parameters given the training data. The posterior predictive distribution can be approximated as 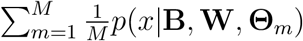 where 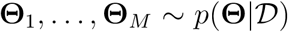 are posterior samples. This distribution can be exactly matched by a stochastic synthesis model using fixed assembly with 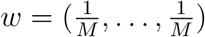 and arbitrary codon mixtures.

#### S2.2 Stochastic EM

We used the online EM algorithm proposed by Cappé and Moulines (2008), modified to update using minibatches instead of individual datapoints. Here we derive the algorithm for the stochastic synthesis model (Equation 1). Without loss of generality, we focus on combinatorial assembly models; the fixed assembly case can be obtained by setting *K* = 1. The local variable of the synthesis model is *Z_i_*, which we represent here as a one-hot encoding, i.e. *Z_i_* ∈ {0,1}^*K*×*M*^. At iteration *t* of the optimization algorithm, given the current parameter estimate *θ*^(*t*)^ = (*w*^(*t*)^, *u*^(*t*)^, *v*^(*t*)^, *τ* ^(*t*)^), the conditional expectation of *Z_i_* can be written as

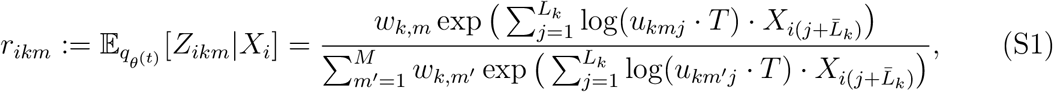

where 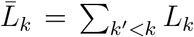. Now we can compute the conditional expectation of the mean log likelihood as

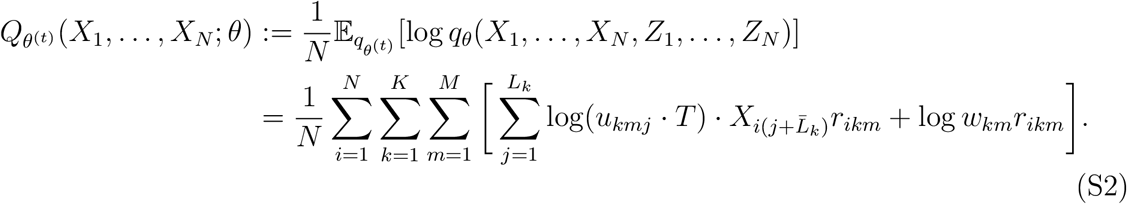

In standard EM, we would optimize this function with respect to *θ.* However, this requires summing over the whole dataset at each step. To derive the stochastic EM algorithm, we rewrite *Q*_*θ*_^(*t*)^ in terms of summary statistics of the data that can be estimated from minibatches. In particular, let 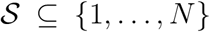 be a subset of the data, and define the summary statistics

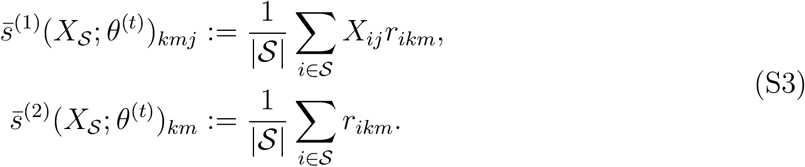

Now we can estimate *Q_θ_*(*t*) as

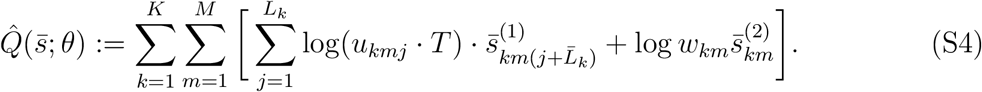

The complete algorithm alternates between estimating summary statistics from minibatches of data 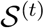 drawn at each step and maximizing the estimated expected log likelihood 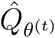,

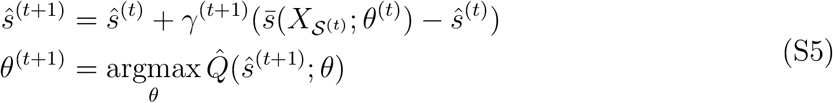

where *γ*^(*t*)^ is the step size. As suggested by Cappé and Moulines (2008), we set *γ*^(*t*)^ = *t*^-0.6^. We also use Polyak-Ruppert averaging, as suggested by Cappé and Moulines (2008), taking the mean of the summary statistics *ŝ*^(*t*)^ for the last half of training, i.e. 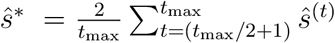, and producing the final parameter estimate 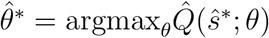.

The maximization step *θ*^(*t*+1)^ = argmax 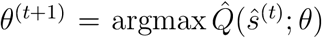 can vary depending on the codon diversification technology used. For all technologies, we have

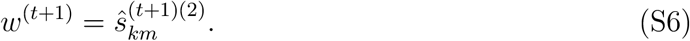

For arbitrary codon mixtures and finite codon mixtures, we can without loss of generality pick one codon for each amino acid and the stop symbol, and work with template probabilities 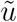 directly over amino acids, i.e. where 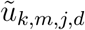 is the probability of amino acid *d* at position *j* of template *m* in pool *k*. Then, for arbitrary codon mixtures,

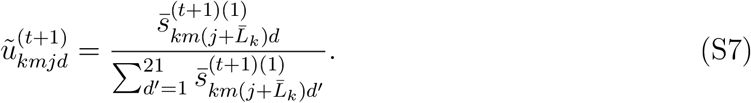

For finite codon mixtures, let 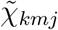 be a one-hot encoding of the codon mixture used at position *j* of template *m* in pool *k*, such that 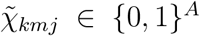. We work directly with mixtures defined over amino acids, with 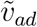 the probability of amino acid *d* in mixture *a*. Thus 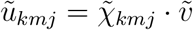. Then we can use the coordinate-wise update

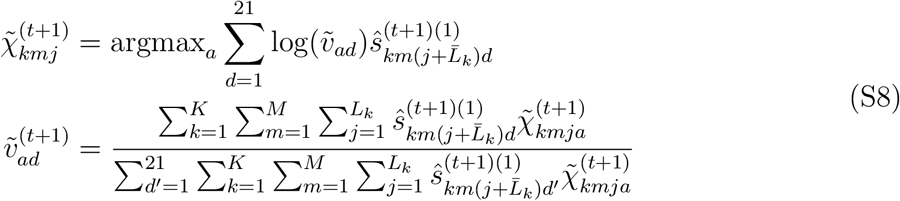

For finite nucleotide mixtures, we use *χ*_*kmj*1_ to denote a one-hot encoding of the mixture used at the first position of the codon at position *j* in template *m* in pool *k*, i.e. *χ*_*kmj*1_ ∈ {0,1}^*A*^, and likewise for *χ*_*kmj*2_ and *χ*_*kmj*3_. We update *χ* by optimizing over all three positions of each codon jointly, enumerating all combinations of *a*_1_, *a*_2_ and *a*_3_,

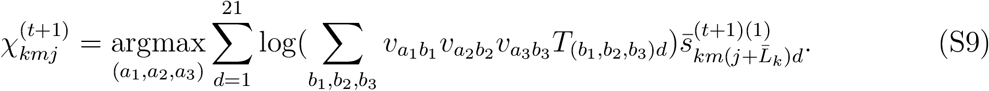

Once *χ* has been updated, we update *v*. This is harder, as there is no closed form solution. We directly optimize 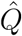 with respect to v by taking gradients and applying 5 steps of the Adam optimizer (Kingma and Ba, 2015) with a learning rate of 0.01 (that is, we take 5 steps of Adam for every 1 EM update). For enzymatic mutagenesis, we can also apply Equation S9 to update *χ*, replacing *v* with *S^τ^*. To update *τ*, we directly enumerate all values of 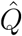 for *τ* ∈ {1,…, *τ*_max_} and choose the maximum.

Code implementing the stochastic EM algorithm for all of the proposed stochastic synthesis models is available in the Supplementary Material.

#### S2.3 Choosing 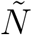

Recall that our proposed black-box optimization procedure is to draw 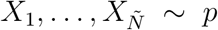 computationally and then maximize the synthesis model parameters,

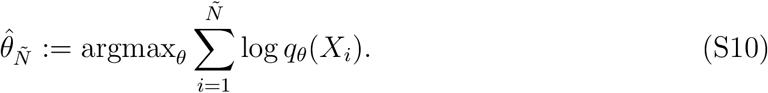

In this section, we argue that 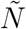 should be chosen to be either equal to *N*_1_, or, if *N*_1_ is too large to be tractable computationally, 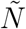 should be as large as is tractable. In particular, we *do not* suggest choosing 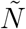 to be larger than *N*_1_, nor do we suggest regularizing *θ* as one would in a standard inference problem. The reason is that “overfitting” the synthesis model to the samples 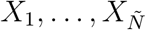 can help rather than hurt.

To be more precise, consider the extreme case where *q_θ_* can exactly match the empirical distribution of *X*_1_,…, *X*_*N*_1__ ~ *p* but cannot exactly match *p* itself. For example, this situation can occur when using fixed assembly and *M* = *N*_1_, allowing each mixture component be a point mass. If we use *N* = *N*_1_, we find

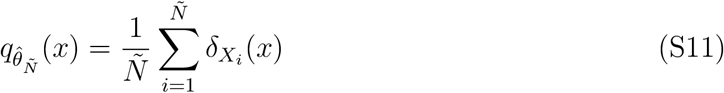

where *δ_x_*′(*x*) is the Kronecker delta function at *x*′. In this case, variational synthesis is equivalent to large-scale MC synthesis, and will produce *N*_1_ samples from *p*.^1^ On the other hand, if we let 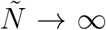, we have 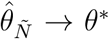. In this case, variational synthesis will produce *N*_1_ samples from *q_θ_\** ≠ *p*. Thus, it can be preferable to use 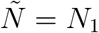 as compared to 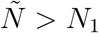, since using 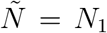 leads to synthesis of *N*_1_ exact samples from *p* instead of *N*_1_ samples from *q_θ*_* ≠ *p*.

In practice, of course, *q_θ_* will rarely be able to exactly match the empirical distribution of samples from *p*. Nonetheless, we expect using 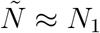 to be useful, as in this case we avoid trying to match 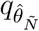 to components of *p* that are too rare to occur in practice, and instead regularize 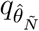 towards the empirical distribution of samples from *p*.

#### S2.4 Variable Length Protein Sequences

To handle variable length protein sequences, we treat everything past the stop codon as missing data which does not contribute to the likelihood. That is, for a sequence *X_i_* with a stop codon at position **j**, we have *q_θ_*(*X_i_*) = *q_θ_*(*X*_*i*,1:**j**_).

### S3 RELATED WORK DETAILS

#### S3.1 DeCoDe

DeCoDe can be applied to datasets of fixed-length (or aligned) sequences, 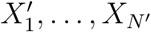, which are assumed to be unique (i.e. 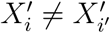, if *i* ≠ *i*′). Consider the empirical distribution 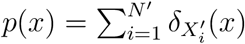 where *δ_x′_*(*x*) is the Kronecker delta. Take *q_θ_* to be a stochastic synthesis model using finite nucleotide mixtures and fixed assembly, with *θ* = (*w, u, v*). Let supp(*p*) denote the support of *p*, i.e. the set of all length *L* sequences with non-zero probability. Let 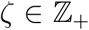 denote the maximum allowed support of *q_θ_*. Then, we can rewrite the DeCoDe objective (Section 2.2.2 in Shimko et al. (2020)) in terms of the size of the intersection of supports of *p* and *q_θ_*,

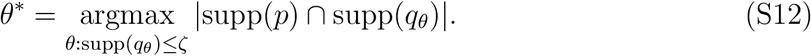

Note that the size of the intersection of supports does not correspond to a valid divergence between *p* and *q_θ_*.

#### S3.2 SCHEMA

RASPP (Endelman et al., 2004) is an algorithm for designing site-directed recombination or combinatorial assembly libraries based on a crystal structure and a dataset of homologous proteins from the same family. It chooses a set of template lengths *L*_1_,…, *L_K_*, where *L*_min_ ≤ *L_k_* ≤ *L*_max_ for *k* ∈ {1,…,*K*}, in order to minimize the SCHEMA score, roughly the number of structural contacts between positions of the protein generated by different template pools. In this section we give a heuristic argument connecting RASPP to variational synthesis, in the special case where RASPP finds a solution with no structural contacts across regions covered by each pool.

Consider a target model *p* that consists of a Potts model learned from the same protein family as the dataset of homologous proteins. In general, the Potts model will infer energetic interactions only between positions of the alignment that are in structural contact (Marks et al., 2011). Let 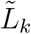 denote the region generated by template *k*, i.e. 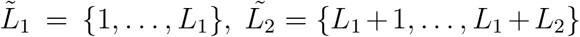, etc. and let 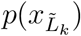 denote the marginal of *p* over these positions. For the set of *L*_1_,…, *L_k_* chosen by RASPP, we have no structural contacts across regions, and so no energetic interactions under the Potts model, and thus 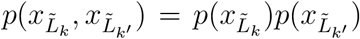 for *k* = *k*^′^. In other words, there is no correlation between segments under the Potts model *p*. When using stochastic synthesis with combinatorial assembly, there is also no correlation between segments under *q_θ_*. If we try to minimize the KL divergence between a *q_θ_* with combinatorial assembly and the Potts model *p*, and optimize the template lengths *L*_1_,…, *L_K_*, we can expect in general to find a similar solution to RASPP, where both the SCHEMA score and the correlation between templates under *p* is zero.

### S4 THEORY DETAILS

Note that the proofs in this section rely on the definitions in Table S1.

#### S4.1 The MC Synthesis Estimator

In our theoretical analysis we do not treat MC synthesis as variational synthesis with point mass (deterministic) mixture components. In particular, we analyze the estimator

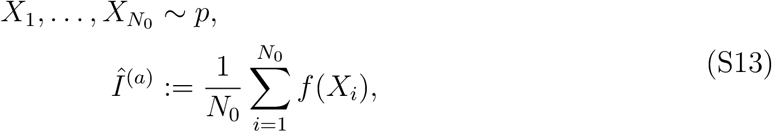

which comes from measuring each synthesized sequence individually, and *not* the alternative estimator

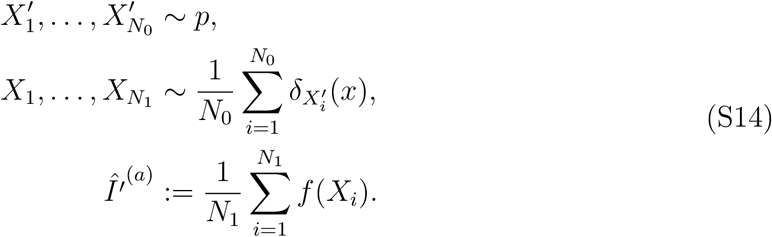

which would come from pooling the synthesized sequences and then measuring a random sample of size *N*_1_ (here *δ_x_*′(*x*) is the Kronecker delta function at *x*′). Note this alternative estimator *Î*′(*a*) takes the form of a bootstrap estimator of size *N*_1_, taken from an initial sample of size *N*_0_ from *p*, and thus in general introduces additional sampling noise as compared to *Î*(*a*). There are three reasons for focusing our analysis on *Î*(*a*) instead of *Î*′(*a*). First, since *N*_0_ is low, in practice it is often tractable for experimentalists to measure the *N*_0_ sequences individually (e.g. in 96 well plates), rather than pooling them, making the estimate *Î*(*a*) possible. Second, in the limit where *N*_1_ is much greater than *N*_0_, the estimators converge, making *Î*(*a*) a reasonable approximation for pooled experiments in practice. Third, we want our analysis to be conservative in measuring the benefits of variational synthesis vis-à-vis the alternative, MC synthesis, so we use the better estimator *Î*(*a*).

#### S4.2 Proof of Proposition 4.1

##### Proof.

Using Jensen’s inequality,

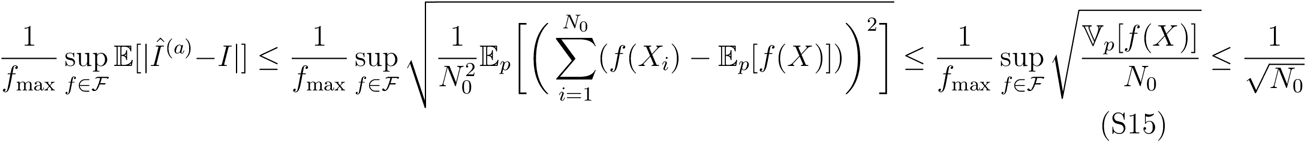

where 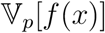 is the variance with respect to *p*.

We can decompose the error in the *Î*(*b*) estimate into variance and bias terms, and then apply a similar analysis,

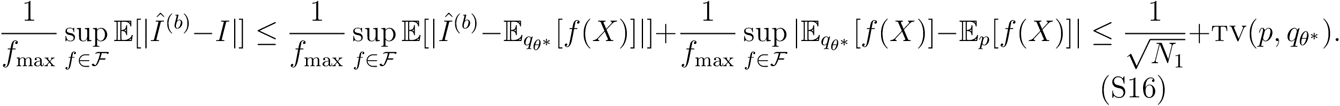

where we have used the integral probability metric representation of the total variation metric tv(·, ·) (Sriperumbudur et al., 2009). The result follows from application of Pinsker’s inequality.

We can see from the proof that the bound in Equation 3 could be tighter if we use total variation in place of KL. It could also be tighter if we restrict the family of functions 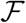 further. In particular, consider the metric space defined over the set of fixed length discrete sequences 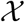 with the Hamming distance 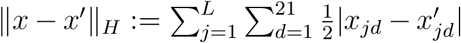 (where x is a one hot encoding of a length *L* nucleotide sequence). Then, we can introduce the function family 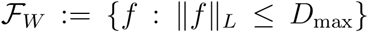, that is, the set of functions with bounded Lipschitz constant 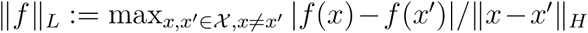. Biologically, the Lipschitz constant is interpretable as the sensitivity of a sequence’s biological function to single mutations. In particular, if a point mutation can dramatically change the assayed property of the sequence, then the Lipschitz constant will be large; otherwise it will be small. If we assume the Lipschitz constant of the experimental assay is bounded by some constant *D*_max_, we can find an alternative error bound on the stochastic synthesis estimator:

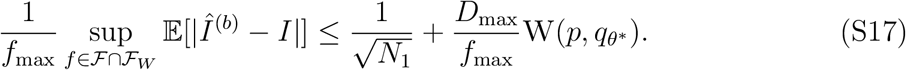

where 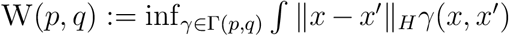 is the first Wasserstein distance, with Γ(*p, q*) the set of couplings of *p* and *q*. This result follows from Equation S16 by applying the Kantorovich-Rubinstein duality theorem (e.g. Dudley (2002), Theorem 11.8.2), using the fact that the metric space of finite sequences with the Hadamard distance is a finite discrete space and separable. We see from Equation S17 that the error bound on variational synthesis can be lower than that in Equation S16, so long as *D*_max_ is sufficiently small. In other words, we can get away with using synthesis models that do not match *p* closely if the assay is not very sensitive to small changes in sequence.

#### S4.3 Importance Sampling Estimates

In some cases we can get access to paired sequence and function data, and in particular the dataset 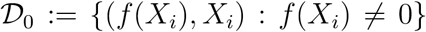. For instance, if we deep sequence the hits of a screen, with 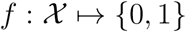, we will have 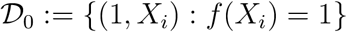. We can then construct an importance-sampling estimate of 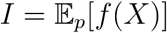 using samples *X*_1_,…, *X*_*N*_1__ ~ *q_θ*_*,

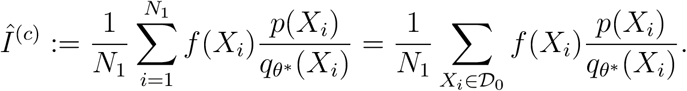

Unlike *Î*(*b*). this estimator is unbiased: 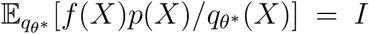. However, *Î*(*c*) still takes advantage of a large number of samples, making possible lower variance than *Î*(*a*). In particular, we have

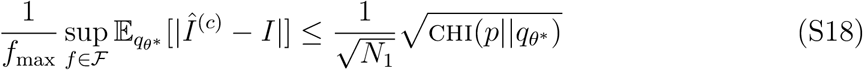

where chi is the chi divergence, which can be defined as 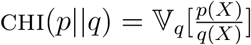. We can derive this result following the same analysis as in Equation S15,

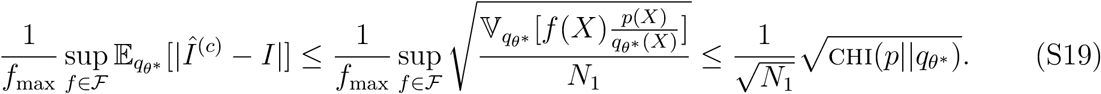

Note that our suggested black-box optimization procedure for variational synthesis (Section 2.2) is intended to help ensure high discovery rates (maximizing 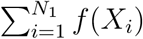) but not to ensure accurate importance sampling estimates. In particular, the kl divergence does not provide a particularly tight bound on the chi divergence (see e.g. Proposition 2 in Dragomir (1999)), so it is likely preferable to (if possible) directly optimize the chi divergence (Dieng et al., 2017).

#### S4.4 Proof of Corollary 4.2

##### Proof.

We have

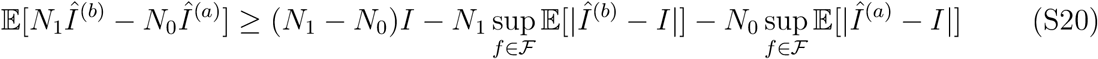

Applying Proposition 4.1 yields the result.

#### S4.5 Proof of Proposition 4.3

Before proving Proposition 4.3, we first prove a lemma that shows – as long as we are not using enzymatic mutagenesis – that we can construct templates that are arbitrarily close to a point mass while still having full support. We use *q_θ_*(*x*|*c*) as shorthand for *q_θ_*(*x*|*C_i_* = *c*), and *δ_x′_*(*x*) to denote the Kronecker delta function which takes value 1 if *x* = *x*′ and 0 otherwise.

##### Lemma S4.1.

*Assume we are using **arbitrary codon mixtures**, **finite codon mixtures** (with A* ≥ 21), *or **finite nucleotide mixtures** (with A* ≥ 4). *For any ϵ* > 0 *sufficiently small, there exists some v such that*:
*for all* 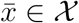 *there exists a* 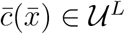 *such that*

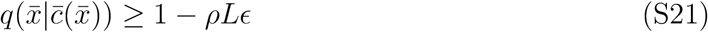

*where ρ is a positive constant, and* 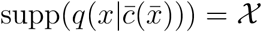. *In particular, for arbitrary or finite codon mixtures, ρ* =1, *while for finite nucleotide mixtures, ρ* = 3.

##### Proof.

We start with the finite codon mixtures case; note that this immediately implies the arbitrary codon mixture case, since the space 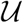 for finite codon mixtures is a subset of the space 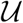 for arbitrary codon mixtures. We choose (arbitrarily) one codon for each amino acid and the stop symbol, and work with mixtures v over these 21 codons (setting the probability of all others to zero). For all *d* ∈ {1,…, 21}, let 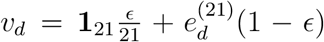 where **1**_*D*_ is the length *D* vector of all ones and 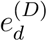 is the length *D* vector of all zeros except a one at position *d*. Let 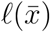 be the length of a protein sequence 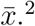 Given 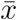 we define the *L* × 21 matrix 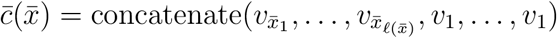. Now note that

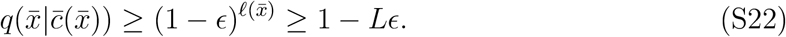

Next we consider the finite nucleotide mixtures case, which works similarly. For all *b* ∈ {1,…, 4}, let 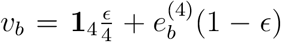. Given a protein sequence 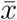, choose a particular codon for each amino acid and the stop symbol. This defines a DNA sequence 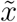, where 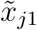 is the nucleotide in the first position of the codon for the amino acid at position *j* of 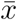, and likewise for 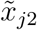 and 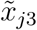. We can then choose nucleotide mixtures for each position of a template to match 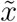, that is,

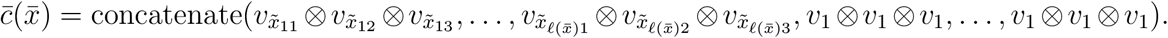

Now we have

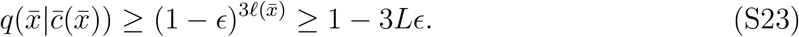

We are now ready to prove Part 1 of Proposition 4.3. The basic idea is to construct a synthesis distribution *q_θ_\** that closely approximates *p* by convolving with *p* templates that are approximate point masses.

#### Part 1 of Proposition 4.3

*When using either **arbitrary codon mixtures**, **finite codon mixtures** (with A* ≥ 21), *or **finite nucleotide mixtures** (with A* ≥ 4): *for any* 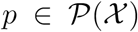 *and η* > 0 *there exists some *M* and θ such that (1)* kl(*p*||*q_θ_*) < *η and (2)* 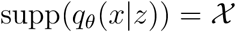 *for all z* ∈ {1,…,*M*}.

*Proof.* Let 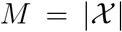, that is, set the number of templates equal to the total number of sequences of length less than or equal to *L*. Since 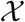 is finite, we can construct for any *ϵ* > 0 the synthesis distribution 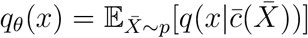. In this synthesis distribution, the weights *w* of each mixture component are set by *p*(*x*), and 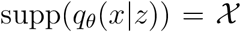 for all *z* by the construction of 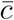. We now have, applying Lemma S4.1,

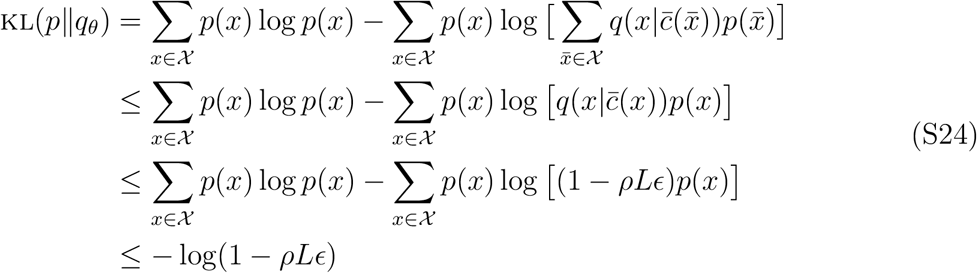

Thus we can choose e sufficiently small that kl(*p*||*q_θ_*) < *η*.

One concerning aspect of this proof, practically, is that it requires very large *M* to form the approximation *q_θ_*. How well can we do with smaller *M*? Combining Theorem 4.2 of Zhang (2003) with the above result, we can say that for any *η* > 0 there exists an *ϵ* > 0 such that kl(*p*||*q_θ*_*) converges to a value less than *η* at a 1/*M* rate. Note also that our setup differs from the more common case where a mixture model is used for density estimation based on finite data, since we can sample as much as we want from *p*. We therefore do not analyze the mismatch between a target *p* and model q_θ_ that may be caused by finite data.

Next, we prove the second part of Proposition 4.3, showing that enzymatic mutagenesis can fail to approximate arbitrary targets *p*. The basic idea is that when using enzymatic mutagenesis, the probability of a particular sequence cannot get arbitrarily close to 1, and so the KL divergence between *p* and *q_θ_* cannot get arbitrarily close to 0.

#### Part 2 of Proposition 4.3

*When using **enzymatic mutagenesis**: there exists some* 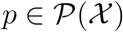 *and η* > 0 *such that for all *M* and θ, we have* kl(*p*||*q_θ_*) > *η.*

*Proof.* Since *τ* > 0, and the entries of *S* are all positive, we can see that we are limited in how much mass an enzymatic mutagenesis model can concentrate on just one sequence, i.e.

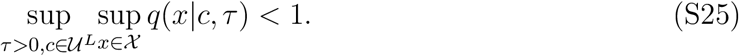

Choose *p*(*x*) = *δ_x′_*(*x*) for some sequence 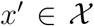, and let *q_θ_* be an enzymatic mutagenesis synthesis model with *M* templates. Then,

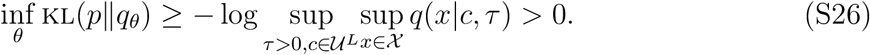

#### S4.6 Proof of Proposition 4.4

##### Proof.

Let 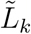 denote the subset of positions generated by template *k*, i.e. 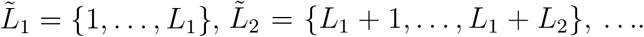 Let 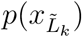 denote the marginal of *p* over these positions. We have, since templates are drawn independently from each pool, 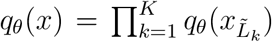, and so

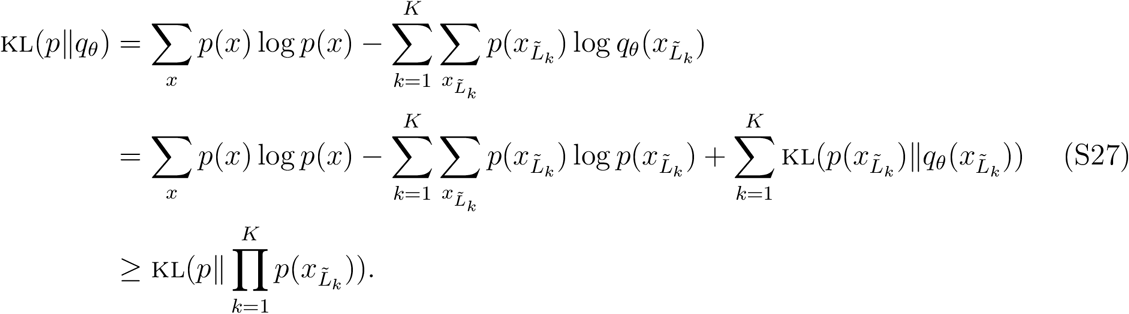

There exists *p* for which 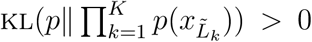, in particular any *p* for which there is correlation between templates, proving the result.

### S5 RESULTS DETAILS

#### S5.1 Datasets and Target Models

##### S5.1.1 DHFR

We used a dataset of 3,629 sequences in the DHFR family collected using jackhmmer (Eddy, 2011) from the Uniref100 dataset (Suzek et al., 2015), and available as an example dataset from https://github.com/debbiemarkslab/plmc/tree/master/example/protein/DHFR.a2m. The multiple sequence alignment has a width of *L* = 171 amino acids. We trained a Potts model using pseuodolikelihood maximization as in Hopf et al. (2017), using the plmc package with the default hyperparameters https://github.com/debbiemarkslab/plmc. Gaps in the alignment were treated as missing data (not as separate symbols), following the default settings of plmc. The trained Potts model was the target *p*. We sampled sequences from *p* using Gibbs sampling, drawing 100,000 samples using 10 parallel chains with a burn-in of 200 steps per chain.

For the analysis of unaligned sequences (Figures 3D and S5), we used the training dataset of 3,629 evolutionary sequences, with gap symbols excluded and stop symbols appended. We refer to this dataset as “DHFR raw”.

##### S5.1.2 GFP

We constructed a dataset of 722 sequences in the GFP family using jackhmmer and UniprotKB (07/2021) (Potter et al., 2018), starting from the seed sequence `GFP_AEQVI` with F64L (a stabilized variant used by Sarkisyan et al. (2016)), with a threshold of 0.3 bit score per residue. We trained an ICA model with a MuE output (Weinstein and Marks, 2021), which is available as an example in the Pyro probabilistic programming language (Bingham et al., 2019) at https://pyro.ai/examples/mue_factor.html. The ICA model is similar to a probabilistic PCA model, but uses a Laplace prior on the latent variable instead of a Gaussian; the MuE output uses the default profile HMM-based architecture described in Weinstein and Marks (2021). We used 2 latent dimensions in the ICA model, a latent sequence length of 237 in the MuE, and default priors. The model was trained with stochastic variational inference, with a learning rate of 0.005 and batch size of 5 over 70 epochs, annealing the prior KL divergence linearly over 35 epochs. Using 20% of the data as heldout validation, the model achieves a per residue perplexity on the training set of 3.1 and on the test set of 4.6.

We used the ICA-MuE model to construct a target distribution *p*. In particular, let *ψ* be the latent alignment variable of the MuE (the state variable of the Markov chain). We estimated the maximum *a posteriori* value of *ψ* for the stabilized wild-type GFP (`GFP_AEQVI` with F64L), and then sampled new sequences conditional on this value 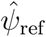 – note that this procedure is a very weak form of supervision, since the stabilized wild-type is known to be functional and produce fluorescence. To limit the diversity of the library relative to the training data, we sampled from the posterior predictive over the latent representation given the observed data, rather than the prior. Explicitly, let *p*_MuE_(*x*|*ψ, κ*) denote the distribution of the learned ICA-MuE model conditional on the latent alignment *ψ* and latent representation *κ*. Let 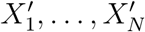, denote the training data, and let *p*(*κ*|*X*′) denote the posterior over the latent representation of a datapoint *X*′ (which can be approximated by the encoder/guide network). The complete generative process *p* is then defined as

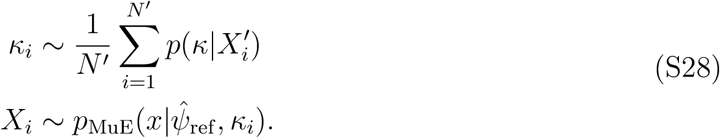

An important feature of this model is that we are *not* sampling from the conditional distribution of *κ* given 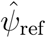, that is, we are not sampling sequences with similar latent alignments. Unlike autoregressive models, for example, MuE models allow variation in sequence length and latent alignment to be treated as independent of variation at conserved sites. Thus, although the sequences generated from *p* are all of the same length, the pattern of amino acids at conserved sites reflects the full diversity of the dataset. Finally, note that in the jackhmmmer constructed-dataset, the first residue (M) of the wild-type sequence `GFP_AEQVI` was not included in the profile HMM envelope, but the sequence-to-function predictor expects this position to be included; we therefore prepended an *M* to each generated sequence, for a total length of *L* = 238.

##### S5.1.3 TCR

We examined a dataset of 22,004 TCR*β* sequences measured in Ramien et al. (2019), taken from CD8+ T cells from a single healthy control patient (number HC12 in the study) in the 3rd trimester of pregnancy. We trained a ICA-MuE model as described above (Section S5.1.2), with 5 latent dimensions and a latent sequence length of 170. We used stochastic variational inference, with a learning rate of 0.01 and batch size of 5 over 2 epochs, annealing the prior KL divergence linearly over 1 epoch. Using 20% of the data as heldout validation, the model achieves a per residue perplexity of 2.39 on both the training and test datasets. We sampled from the model using the same strategy as in Section S5.1.2. The reference sequence used to construct 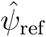 was a randomly selected sequence from the dataset described in Section S5.6.2, which Tcellmatch predicted to bind the influenza epitope; as with the GFP example, conditioning on 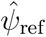 is a very weak form of supervision, learning from only a single functional example. In particular, the reference sequence was *MSNQVLCCVVLCLLGANTVDGGITQSPKYLFRKEGQNVTLSCEQNLNHDAMYWY RQDPGQGLRL/YYSQ/VNDFQKGD/AEGYSVSREKKESFPLTVTSAQKNPTAFYLC ASSZRSAYEQYFGPGTRLTVTEDLKNVFPPEVAVFEPSE*. The generated sequences had length *L* = 149.

### S5.2 Synthesis Model Hyperparameters

In this section we describe the details of our stochastic synthesis models and optimization procedure. We used *K* = 5 pools, with *L_k_* of approximately the same length for each *k* ∈ {1, …, *K*} (the last template was shortened as necessary since *L* is not always a multiple of 5). This yields templates of length 29 to 48 amino acids across all the datasets considered, which is consistent with typical oligosynthesis lengths of ~ 150 nucleotides. We used *A* = 8 for finite nucleotide mixtures; this value is realistic, as the company IDT, for example, currently offers four custom mixtures per oligo plus preset mixtures and single nucleotides (Pazdernik and Bowersox, 2016). We used *A* = 24 for finite codon mixtures, which is similar to typical trimer-based synthesis projects, which use the 20 amino acids plus a few custom mixtures (McMahon et al., 2018).

We set the mutation matrix S based on the ePCR enzyme Mutazyme II, available as part of Agilent’s GeneMorph II Random Mutagenesis Kit https://www.chem-agilent.com/pdf/strata/200550.pdf. In particular, we converted the reported mutational spectra (Table II) into a substitution matrix, under the assumption that the test sequences are 50% A-T base pairs and 50% G-C base pairs: for instance, the probability of a particular base pair mutating per round of mutagenesis is given as 1% overall (10 bases per kilobase), and 50.7% of mutations happen to A-T base pairs, so the probability of a particular A-T base pair mutating is 0.01 o 0.507/0.5. Proceeding in this way, we find

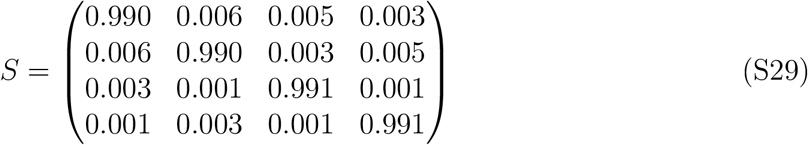

where the columns and rows are each in the order *A, T, G, C*. We also computed a mutation matrix S based on the Taq error prone polymerase (also in Table II of the Gene Morph II Random Mutagenesis Kit manual), but preliminary experiments suggested worse performance than Mutazyme II at matching the DHFR Potts target distribution, so we did not pursue it further. We limit the total number of rounds of mutagenesis *τ* to be less than 10, since large numbers of mutagenesis rounds are rarely used in practice.

Note that since we have chosen *A* ≥ 4, the set of allowed values of 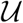 for enzymatic mutagenesis (that is, for all *τ*) is a strict subset of the set of allowed values of 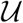 for finite nucleotide mixtures (that is, for all *v*); thus, synthesis models using enzymatic mutagenesis are strictly less expressive than those using finite nucleotide mixtures. Meanwhile, 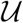 for finite nucleotide or codon mixtures is a strict subset of 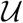 for arbitrary codon mixtures, regardless of the choice of v; so synthesis models using finite mixtures are strictly less expressive than those using arbitrary codon mixtures.

### S5.3 Baseline Synthesis Model

As a baseline stochastic synthesis approach, we considered a method motivated by a common heuristic for producing diversified libraries, which is to simply perform error prone PCR on an initial set of sequences. In particular, the baseline approach we consider is to do MC synthesis plus enzymatic mutagenesis: sample initial protein sequences 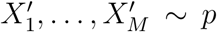, inverse-translate the protein sequences into DNA (sampling uniformly among all codons for the same amino acid), synthesize the DNA individually, and then mutagenize in the laboratory using ePCR. The distribution of resulting sequences can be described using a stochastic synthesis model for which we do *not* optimize the parameters. In particular, let *K* = 1, and for *m* ∈ {1,…,*M*} and *j* ∈ {1,…,*L*}, let 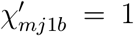 if the sampled codon for 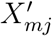 has base *b* at the first position, and *χ*_*mj*1*b*_ = 0 otherwise. Likewise for *χ*_*mj*2*b*_ and *χ*_*mj*3*b*_. Then, we set *u*_1*mj*_ = *S*^*τ*^*χ*_*mj*1_ ⊗ *S*^*τ*^*χ*_*mj*2_ ⊗ *S*^*τ*^*χ*_*mj*3_. We use fixed assembly, setting 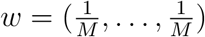. Then, the complete synthesis model (Equation 1) describes the distribution of sequences produced by the baseline approach.

**Figure S1:**
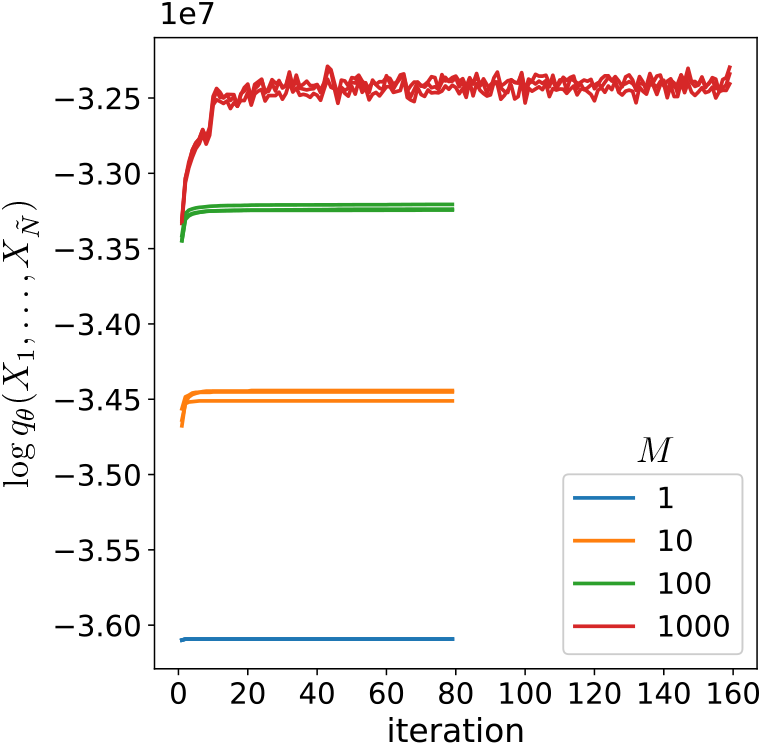
Illustrative example of training curves for a stochastic synthesis model (enzymatic mutagenesis with fixed assembly) with different values of *M*. For each value of *M*, training is repeated with three initial seeds. The models are each trained on samples from the DHFR Potts model, as described in Section S5.4.

Note that the baseline is effectively a kernel density estimate of *p*. It is thus unsurprising that the baseline underperforms relative to variational synthesis, since kernel density estimates typically underperform compared to mixture models.

Practically, we use *S* corresponding to a Mutazyme II enzyme (Section S5.2) and set *τ* = 5 as a typical value for proteins of the length considered here (Wilson and Keefe, 2001). The samples from *p* used as initial sequences, 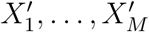, are subsampled from the same training dataset of 100,000 sequences used for variational synthesis (Section S5.4). We examined the performance of the method averaged over 3 independent sets of initial sequences.

### S5.4 Optimization and Perplexity Evaluation

#### DHFR Potts, GFP, TCR

To optimize synthesis models, we drew 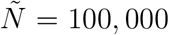 samples from each target distribution *p* and applied stochastic EM, as described in Section S2.2. We chose batch sizes to be as large as possible without running out of memory. In particular, we used batch sizes of 100,000 (the full dataset) with *M* = 1, *M* = 10 and *M* = 100, and batch sizes of 10,000 for *M* = 1000. We trained for 80 epochs with *M* = 1, *M* = 10 and *M* = 100, and 16 epochs for *M* = 1000. Training took 2-5 minutes for each target-synthesis pair using a Tesla V100 GPU. Example training curves are shown in Figure S1.

Each synthesis model was trained on the same set of 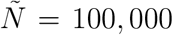 samples from each target distribution, and evaluated based on the average per residue perplexity on the training dataset, 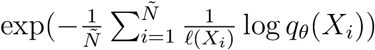, where ℓ(*X_i_*) is the length of the sequence *X_i_*. Note that we do not perform heldout evaluation, as our goal is to see how well each synthesis model can match a target library of size 100,000; overfitting the synthesis model is not a concern, and may even help downstream performance, as described in Section S2.3. We initialized each optimization from three random seeds, and chose the result with the lowest perplexity.

#### DHFR raw

For the DHFR raw dataset, we handle variable length sequences as described in Section S2.4, and optimized each synthesis model using EM with batch size of 3,629 (the full dataset), for 100 epochs. We set *L* to be the maximum length of sequences in the dataset including the stop codon, 170. We evaluated using mean per residue perplexity on the full dataset. We initialized each model from three random seeds, and chose the result with the lowest perplexity.

### S5.5 BEAR Two-Sample Test

We use the vanilla version of the BEAR two-sample test proposed in Section 5 of Amin et al. (2021) to compare the target and the synthesis distributions. The test computes the Bayes factor comparing the hypothesis that two datasets 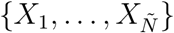 and 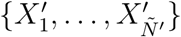 come from the same underlying distribution versus different distributions. It uses 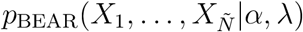, the probability of the dataset under a Bayesian Markov model with Dirichlet concentration parameter *α* and lag λ. In particular, the Bayes factor is

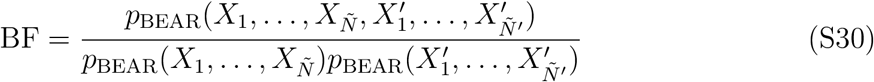

where

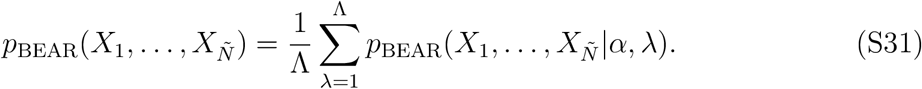

We used the training dataset of 100,000 samples from *p* as the first dataset in the two-sample test, and 100,000 independent samples drawn from the optimized synthesis model 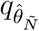 as the second dataset. (In the case of DHFR raw, we used the 3,629 sequences as the target sample.) Note that the goal here is to understand whether the particular set of 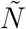 samples from *p* used for training look like a plausible set of samples from 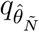, following the logic of Section S2.3, so we do not resample from *p* to compute the test. We use *α* = 0.5 and Λ = 8; we found that in general the posterior over lags concentrated at values of λ below 8, suggesting the test has sufficiently high resolution. Computing the test took about 5-10 minutes for each target-synthesis pair, with 20 cores on an Intel Xeon E5 v3 CPU.

### S5.6 Sequence-to-Function Predictors

#### S5.6.1 GFP: TAPE

We computed TAPE predictions of GFP fluorescence using the interface in the FLEXS package (Sinai et al., 2020). Sequences with internal stop codons were assigned the minimum log fluorescence in the Sarkisyan et al. (2016) dataset, 1.2. Variants with predicted log fluorescence above 3 were classified as hits, in line with the analysis of Sarkisyan et al. (2016) who classify variants below 3 as dark.

#### S5.6.2 TCR: Tcellmatch

We used Tcellmatch, trained on the same single-cell TCR sequencing data as in the original paper (10x Genomics, 2019), with the suggested model architecture (1×1 convolutional embeddings based on BLOSUM50 and biGRU layers). We used the mean squared logarithmic error to evaluate the model’s ability to predict MHC multimer binding counts. We trained the Tcellmatch model only on TCR*β* sequences, since the target *p* was trained only on TCR*β* sequences. The Tcellmatch model uses only the CDR3 region to make predictions. In general, techniques for identifying the CDR3 region in TCRs rely on nucleotide-level information, which is unavailable for generated amino acid sequences. However, we constructed the target *p* by conditioning on a latent alignment, which in turn is based on a reference sequence with nucleotide-level information (Section S5.1.3). We thus use the positions corresponding to the CDR3 in the reference sequence (109:122, as annotated by the 10x pipeline) to define the CDR3 for each sampled sequence from *p* and *q_θ_*. Although the Tcellmatch model can be used to predict many different antigens, we focused on predictions of the GILGFVFTL influenza antigen, since the model had the most accurate predictions for this particular antigen (according to the *R*^2^ metric used by Fischer et al. (2020), in particular *R*^2^ = 0.60). We conditioned on a single donor (donor 1) when making predictions with Tcellmatch. Sequences with internal stop codons were assigned zero counts. Variants with predicted counts above 10 were classified as hits, in line with the analysis of Fischer et al. (2020).

#### S5.6.3 Estimating Hit Rates

Given a dataset of indicators for whether or not each of 100,000 samples from *q_θ_* was a hit or not, i.e. {*f*(*X*_1_),…, *f*(*X_N_*)} where 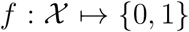, we estimated the overall hit rate using a Beta(0.5, 0.5) prior (Jeffreys prior). We report the standard deviation of the posterior in Figure 4C and G.

#### S5.6.4 Estimating the Number of Unique Hits

Based on the hit rate (Section S5.6.3), we can estimate the total number of hits for libraries of any size. However, we are also interested in the total number of unique hits, since discovering identical sequences is not as useful as discovering diverse sequences. Evaluating predictors on very large numbers of samples, though, can be impractical since predictors (especially TAPE) can be computationally expensive. Instead, we used a Good-Toulmin estimation strategy: we examined the hits from a sample of 100,000 sequences from *q_θ_* and then extrapolated to estimate the number of unique hits in a library of 1,000,000 sequences. We used the smoothed Good-Toulmin estimator proposed by Orlitsky et al. (2016), with the recommended Binomial model. Note that the estimator is considered trustworthy for datasets up to a factor of log *N* larger than the initial dataset; since log(10^5^) = 11.5 ≥ 10, it is applicable here. We estimate the variance of the estimate under resampling using the jackknife, which can be efficiently computed for the smoothed Good-Toulmin estimator (Efron and Stein, 1981).

**Figure S2:**
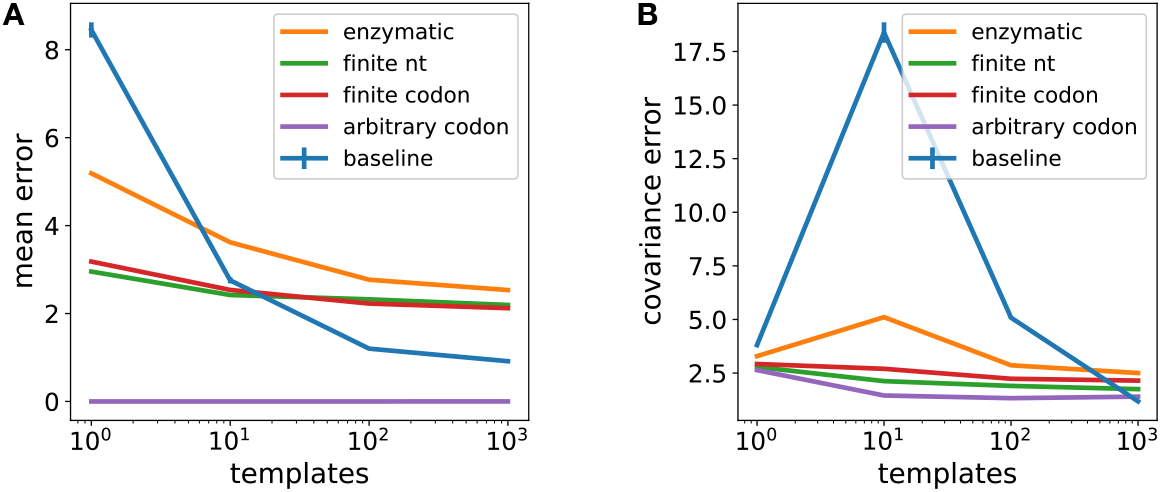
(A) Difference in mean between synthesis and target models, for various stochastic synthesis models with fixed assembly applied to the DHFR Potts target. Mathematically, 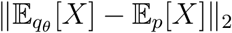 where *X* is represented as a one-hot encoding and || · ||_2_ is the Euclidean distance. (B) Difference in position-wise covariance matrices between synthesis and target models. Mathematically, let 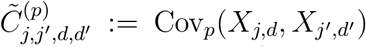 denote the covariance under *p* between the *d*th amino acid at position *j* and the *d*′th amino acid at position *j*′. The magnitude of the covariance between positions *j* and *j*′ can be measured as 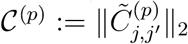. Then we plot the position-wise covariance error 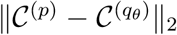. In both plots, error bars for the baseline model are the standard deviation over initial sequences (Section S5.7).

### S5.7 Error Bars

In this section we summarize the calculation of the error bars in Figures 3 and 4. For the baseline model, we show the estimated standard deviation across independent samples of the initial *M* sequences from *p* (that is, the initial sequences that are mutagenized by ePCR). We use three independent samples for each value of *M*. For perplexity plots (Figures 3ACD and 4AE) we do not include any error estimates for non-baseline models, since we have exactly computed the total perplexity across the training dataset, and we are only interested in the match between the synthesis model and the training dataset, not in the synthesis model’s generalization performance (as explained in Section S2.3). Bayes factors are themselves measurements of statistical significance, so we do not include any error bars for non-baseline models in Figures 3B and 4BF. For plots of hit rate (Figure 4CG), error bars show the posterior standard deviation of the hit rate under a Beta(0.5, 0.5) prior (the Jeffreys prior) (Section S5.6.3). For plots of estimated unique hits (Figures 4DH), error bars show the jackknife estimate of the standard deviation (Section S5.6.4) For the baseline model, in plots of both hit rate and unique hits, error bars include the variance across different initial sequences from *p*, and are computed using the law of total variance.

### S5.8 Additional results

#### S5.8.1 DHFR

We further examined the match between stochastic synthesis models and the target DHFR Potts model, examining the difference in moments of each distribution. In particular, we looked at the difference in the mean sequence produced by the synthesis and target distributions, and the difference in covariance between positions of the sequences produced by the synthesis and target distributions (Figure S2). Comparing different variational synthesis models, we see improved perplexity (Figure 3A) corresponds well with lower moment error (Figure S2). Interestingly, the baseline synthesis method (Section S5.3) yields comparatively low moment error for large *M* despite comparatively poor perplexity.

**Figure S3:**
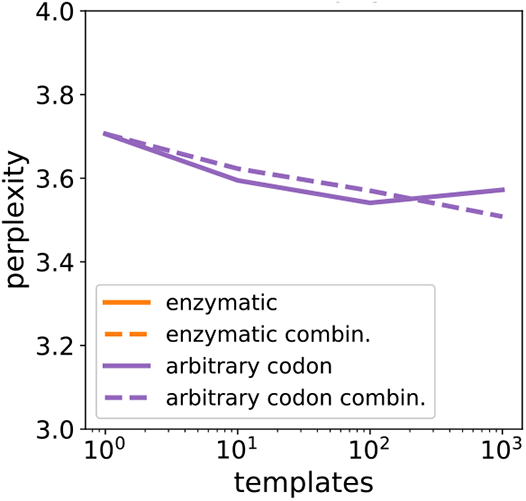
Zoom in of Figure 3C.

**Figure S4:**
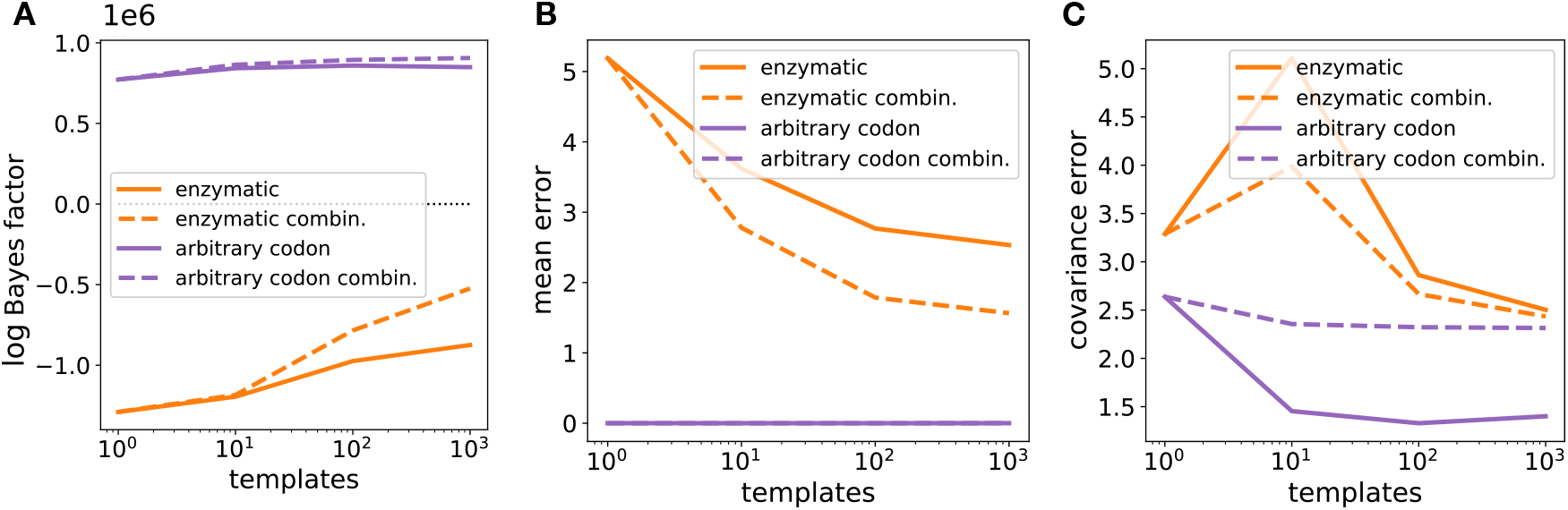
Comparing fixed versus combinatorial assembly for the DHFR Potts target. (The perplexity comparison can be found in Figure 3C and S3.) (A) Two-sample test Bayes factor. (B) Difference in mean between synthesis and target models, as defined in Figure S2. (C) Difference in position-wise covariance matrices between synthesis and target models, as defined in Figure S2.

We further examined the difference in performance between combinatorial and fixed assembly models. For enzymatic mutagenesis, switching from fixed to combinatorial assembly improves the two-sample test Bayes factor (Figure S4A), mean (Figure S4B) and covariance (Figure S4C). For arbitrary codon synthesis, switching from fixed to combinatorial assembly slightly improves the Bayes factor (Figure S4A), has no effect on the mean (as we expect mathematically and see in Figure S4B), but substantially worsens the covariance (Figure S4C). These results illustrate how the advantages of using fixed versus combinatorial assembly vary depending on the codon diversification technology.

We further examined the performance of different stochastic synthesis models applied to the DHFR raw dataset of unaligned evolutionary sequences. Applying the two-sample test, we find that using large numbers of templates with any codon diversification technology is better than using small numbers of templates with a very expressive codon diversification technology (Figure S5), in line with the perplexity results (Figure 3D). We also see that variational synthesis is capable of matching the target closely enough to pass the two-sample test, but so is the baseline method in this case.

**Figure S5:**
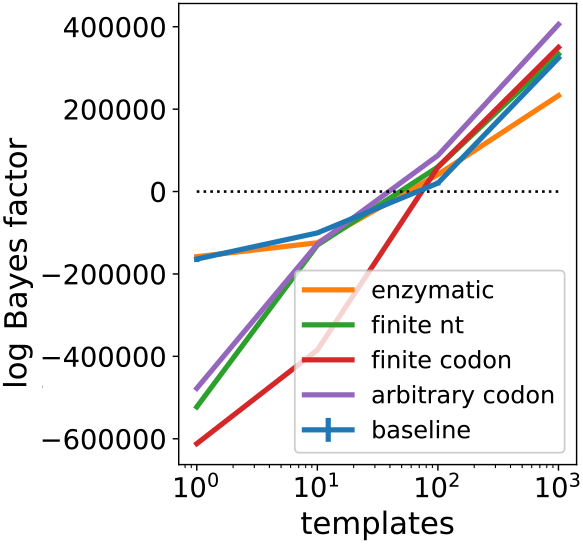
Two-sample test Bayes factor for synthesis models with fixed assembly applied to the DHFR raw dataset. For perplexity comparison, see Figure 3D.

**Figure S6:**
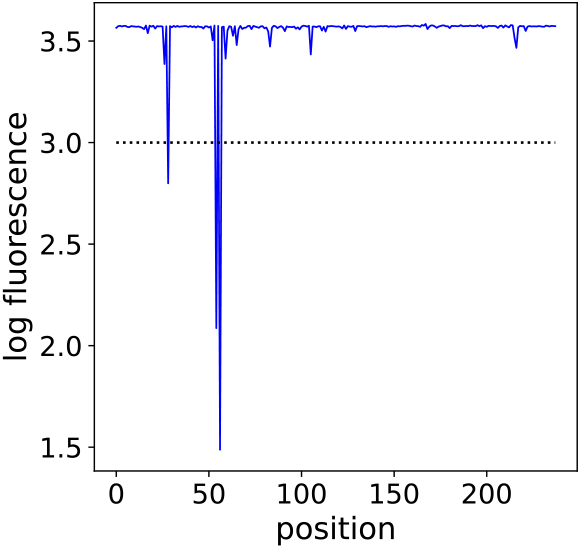
Predicted mutational effects of substituting each position of the stabilized wildtype GFP sequence with an alanine (i.e. an *in silico* alanine scan). Dotted line shows the threshold for classifying a variant as functional.

#### S5.8.2 GFP

We examined the difference in moments between the target GFP distribution and the stochastic synthesis models. The results (Figure S7) are qualitatively similar to those described for DHFR (Section S5.8.1 and Figure S2), with the baseline model performing better than its perplexity would suggest.

We examined the difference in average log fluorescence between samples from various stochastic synthesis models, as compared to exact samples from the target (that is, as compared to the average log fluorescence under MC synthesis) (Figure S8). Interestingly, we find that while using finite codon mixtures with *M* = 1 yields relatively low hit rates compared to arbitrary codon mixtures with *M* = 1 (Figure 4G), it yields nearly equivalent average log fluorescence (Figure S8).

We examined the difference in performance between combinatorial and fixed assembly methods applied to the GFP target distribution. On statistical measures of the difference between the synthesis and target distribution (Figure S9ABCD), we find broadly similar effects to those observed for DHFR Potts: for instance, we see moderate improvements in perplexity for enzymatic mutagenesis at large *M* when switching from fixed to combinatorial assembly, but little effect for arbitrary codon mixtures, and substantially worse covariance for arbitrary codon mixtures. On measures of function, using combinatorial assembly leads to dramatically worse performance (Figure S9EFG): using arbitrary codon mixtures with combinatorial instead of fixed assembly drops the number of unique hits by three orders of magnitude. This result suggests that passing the BEAR two-sample test with large Bayes factors is not enough to ensure high hit rates when using combinatorial assembly; one should also inspect the covariance error.

**Figure S7:**
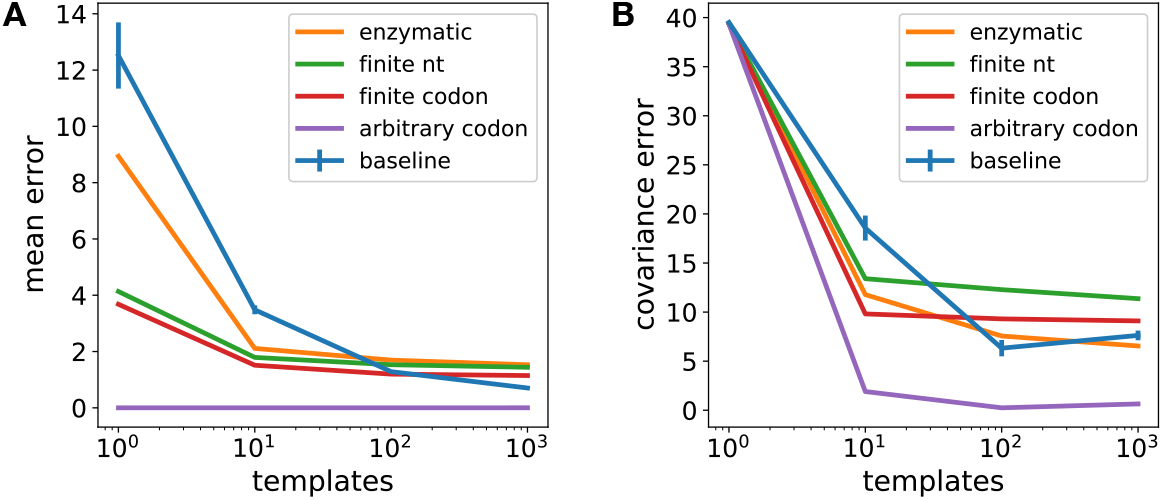
(A) Difference in mean between GFP synthesis and target models, as defined in the caption of Figure S2. (B) Difference in position-wise covariance matrices between GFP synthesis and target models, as defined in the caption of Figure S2.

**Figure S8:**
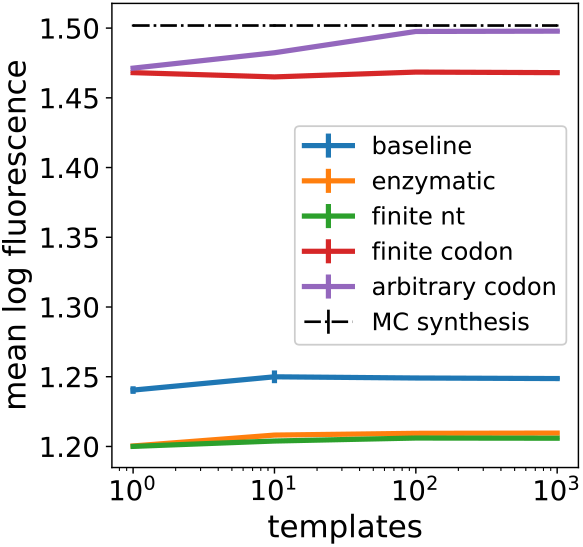
Average log predicted fluorescence of samples from various stochastic synthesis models and from the GFP target *p* itself (MC synthesis). Error bars are estimates of the standard deviation of the mean (for the baseline model, this includes variance across different initial sequences, as described in Section S5.7; for the rest of the models, it is just the standard error, and negligible in these plots).

#### S5.8.3 TCR

We examined the difference in moments between the target TCR distribution and the stochastic synthesis models. The results (Figure S11) are qualitatively similar to those described for DHFR (Section S5.8.1 and Figure S2), with the baseline model performing better than its perplexity would suggest.

**Figure S9:**
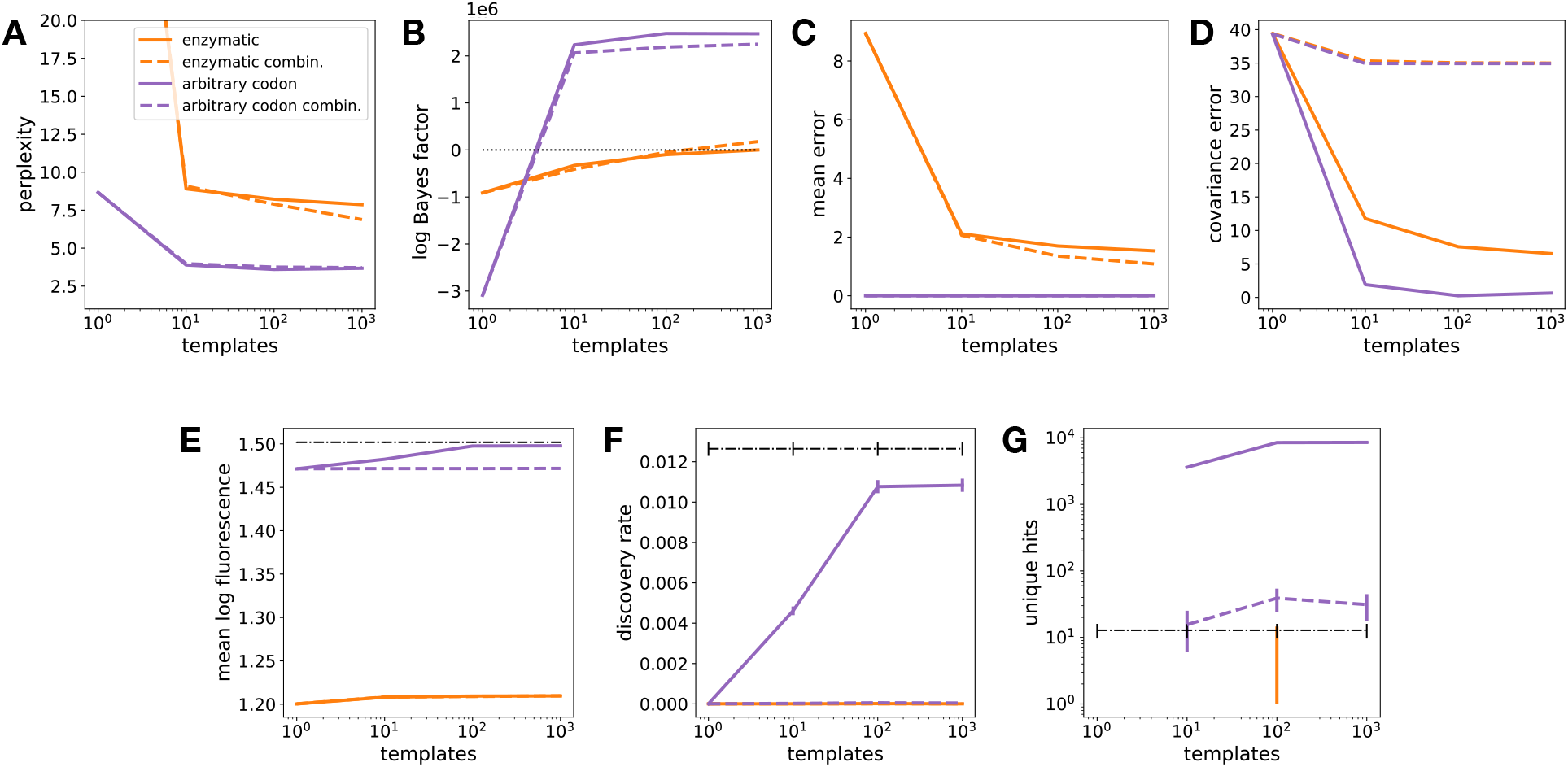
Fixed versus combinatorial stochastic synthesis applied to the GFP target distribution. (A) perplexity, (B) two-sample test Bayes factor, (C) mean error (as defined in Figure S2), (D) covariance error (as defined in Figure S2), (E) average log fluorescence, (F) hit rate and (G) number of unique hits with *N*_1_ = 10^6^ and *N*_0_ = 10^3^. Error bars are as described in Section S5.7.

We examined the difference in average binding counts between samples from various stochastic synthesis models, as compared to exact samples from the target TCR model (that is, as compared to MC synthesis) (Figure S12). Unlike for GFP, we find that the average value of the assay output is roughly proportional to the hit rate.

---

**Figure S10:**
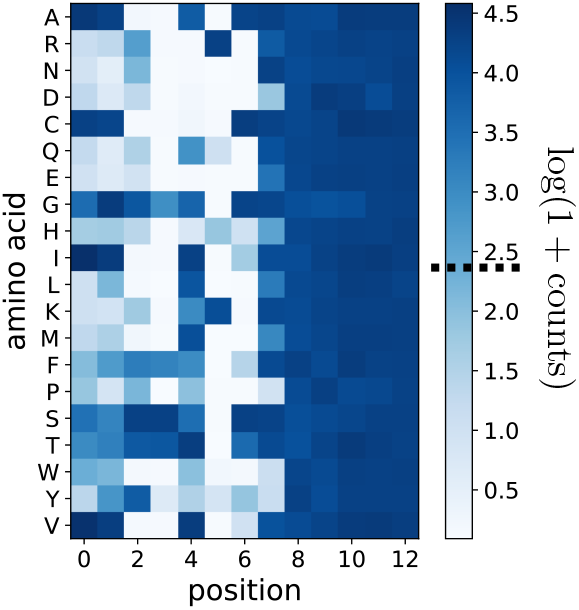
Predicted binding effects of substituting each position of a natural CDR3 sequence (*CASSIRSAY EQY F*) with each of 20 amino acids (*in silico* deep mutational scan). The threshold for functionality (10 counts) is marked by a dotted line in the colorbar.

**Figure S11:**
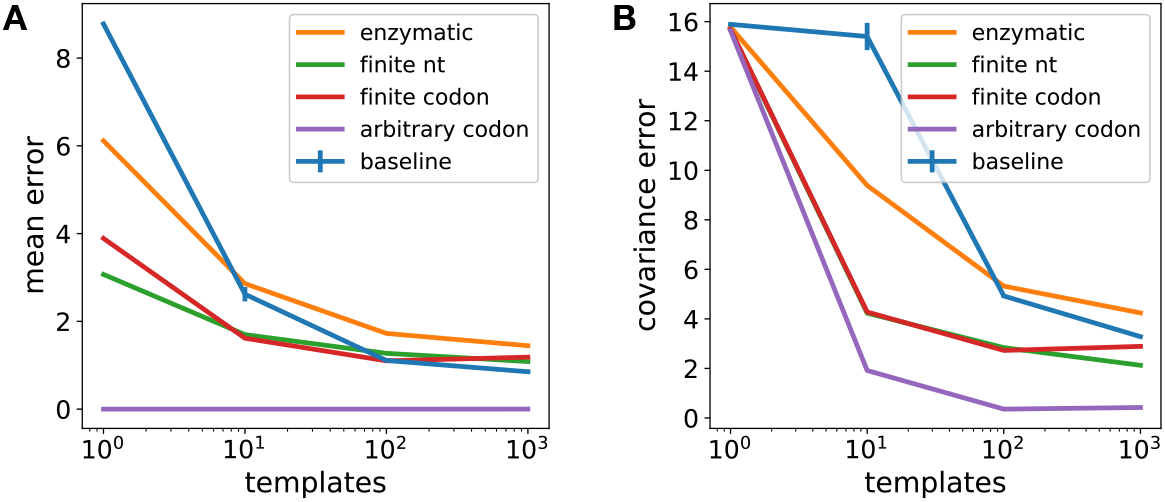
(A) Difference in mean between TCR synthesis and target models, as defined in the caption of Figure S2. (B) Difference in position-wise covariance matrices between TCR synthesis and target models, as defined in the caption of Figure S2.

**Figure S12:**
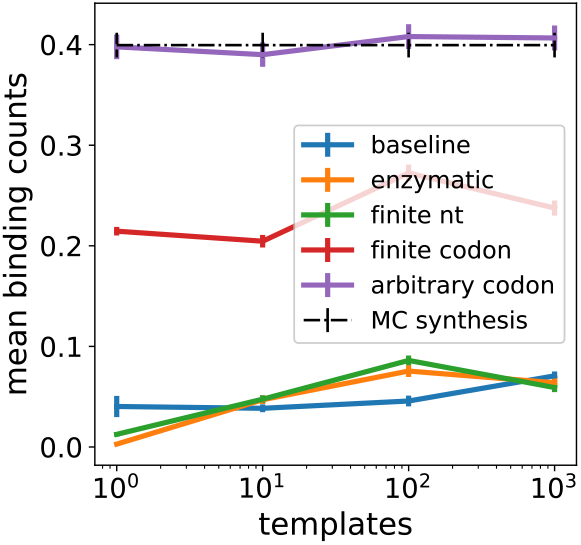
Average predicted binding counts of samples from various stochastic synthesis models and from *p* itself (MC synthesis). Error bars are estimates of the standard deviation of the mean (for the baseline model, this includes variance across different initial sequences; for the rest of the models, it is just the standard error).

## Footnotes

1 We index the 64 codons either using either tuples (*A, A, A*),…, (*T, T, T*) or integers 1,…, 64, depending on convenience.

2 In practice, despite the mathematical idealization of our models, all synthesis technologies have a minimum non-zero codon probability, set by engineering constraints. The key question is really how low this number is comparatively.

1 Technically, variational synthesis in this case produces a size *N*_1_ bootstrap of *N*_1_ samples from *p*, rather than directly producing *N*_1_ samples from *p*. Although bootstrapping introduces some additional sampling noise, we expect it is unlikely in practice to make using 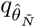 worse than using *q_θ*_*, since the bootstrap directly approximates *p*. Section S4.1 discusses this subtlety further.

2 Length is measured up to (and including) the first stop codon or *L*, whichever comes first.

